# Mechanical strain can increase segment number in live chick embryos

**DOI:** 10.1101/211623

**Authors:** Ben K. A. Nelemans, Manuel Schmitz, Hannan Tahir, Roeland M. H. Merks, Theodoor H. Smit

## Abstract

Physical cues, experienced during early embryonic development, can influence species-specific vertebral numbers. Here we show that mechanical stretching of live chicken embryos can induce the formation of additional somites and thereby modify early segmental patterning. Stretching deforms the somites, and results in a cellular reorganization that forms stable daughter somites. Cells from the somite core thereby undergo mesenchymal-to-epithelial transitions (MET), thus meeting the geometrical demand for more border cells. Using a Cellular Potts Model, we suggest that this MET occurs through lateral induction by the existing epithelial cells. Our results indicate that self-organizing properties of the somitic mesoderm generate phenotypic plasticity that allows it to cope with variations in the mechanical environment. This plasticity may provide a novel mechanism for explaining how vertebral numbers in species may have increased during evolution. Additionally, by preventing the formation of transitional vertebrae, these self-organization qualities of somites may be selectively advantageous.

## Introduction

A segmented spine is the characteristic feature of the vertebrate body plan, which provides mechanical support, flexibility, and protection of the spinal cord. Vertebral numbers vary considerably among species, ranging from six in frogs to several hundred in snakes (Richardson, Allen, Wright, Raynaud, & Hanken, 1998). The evolvability of the vertebral column allows vertebrates to adapt to diverse habitats and acquire matching locomotor styles by tuning the number of body segments (Galis et al., 2014a).

Patterning of the vertebrate body originates early in embryogenesis when the paraxial mesoderm on both sides of the midline segments into somites, which contain the predecessor cells of vertebrae, ribs, muscles and skin. Somites are cell blocks in which a core of mesenchymal cells, the somitocoel, is surrounded by an epithelial layer (Kulesa, Schnell, Rudloff, Baker, & Maini, 2007; Martins et al., 2009); they also impose a segmented organization on the peripheral nervous system. The sequential partitioning of the paraxial mesoderm appears to be imposed by a molecular clock and a travelling wave of maturation created by a system of signalling gradients (Hubaud & Pourquié, 2014). This complex signalling network appears well conserved throughout the vertebrate phylum. The variation in somite numbers (and vertebral numbers) between species is presumably caused by mutations that lead to changes in the speed of the clock period or the elongation rate of the mesoderm, which both affect segmentation rate and somite size (Gomez et al., 2008; Gomez & Pourquie, 2009; Herrgen et al., 2010).

Despite the intricately controlled network of the clock-and-wavefront mechanism, vertebral numbers in fish (Beacham & Murray, 1986; Hubbs, 1922; Tibblin, Berggren, Nordahl, Larsson, & Forsman, 2016), amphibians (Jockusch, 1997; Peabody & Brodie Jr., 1975), reptiles (Osgood, 1978), mammals (Lecyk, 1966) and birds (Lindsey & Moodie, 1967) may be influenced during early embryonic development by environmental cues such as temperature, salinity or light conditions. Mechanics may be another physical cue that induces different phenotypes of segmental patterning. In the framework of the Extended Evolutionary Synthesis, it has been argued according to the ‘side-effect hypothesis’, that morphological novelties in the body plan can result not only from genetic rearrangements, but also from mechanical cues exerted on self-organizing developing tissues (Laland et al., 2015; G. Müller, 1990; G. B. Müller, 2003). Somitic mesoderm possesses a certain capacity for self-organization, as somite-like structures can form ectopically in the absence of a clock or a wavefront (Dias, de Almeida, Belmonte, Glazier, & Stern, 2014). The size of epithelializing somites strongly correlates with embryonic growth (Tam, 1981), and hampering axial elongation of the embryo leads to disorganized somites (Stern & Bellairs, 1984). Somite formation also requires the contraction of cells in the PSM (Duess, Fujiwara, Corcionivoschi, Puri, & Thompson, 2013) and mechanical adhesion to the surrounding fibronectin matrix (Hubaud, Regev, Mahadevan, & Pourquié, 2017; Martins et al., 2009).

On the basis of these observations, we hypothesized that if somitic mesoderm is self-organizing under the influence of biomechanical cues, mechanical stretching may then suffice to induce morphological changes in the segmented vertebrate body plan.

To test this hypothesis, we developed a novel experimental setup to apply controlled strains to live chicken embryos. Time-lapse imaging confirmed that, given sufficient strain, mesodermal patterning can indeed be modulated (Fig S1). This suggests that mechanical cues can affect morphogenesis and may be an explanatory factor for variation in vertebral numbers in the scope of the ‘side-effect hypothesis’.

## Results

To test the effect of mechanical stretching on somitogenesis, stage HH8-9 chick embryos (Hamburger & Hamilton, 1951) were cultured *ex ovo* in modified submerged filter paper sandwiches (Schmitz, Nelemans, & Smit, 2016) and stretched along their body axis twice for 51 to 54 minutes at 1.2 μm/s (Fig S1). As a result, the embryos experienced strain values of 23 ± 3% (average ± SD) after the first pull and 19 ± 3% after the second pull. Approximately 12 hours after manipulation, the mesodermal segmentation of one third (19/57) of all stretched embryos was clearly disturbed, unlike that of control embryos (Fig 1A and B). Time-lapse microscopic imaging (Schmitz et al., 2016) revealed that somites budded off from the PSM at regular intervals of 79 ± 8 minutes in controls and 80 ± 6 minutes in stretched samples. Stretched somites were more elongated than those in control embryos (Fig 1C-F, Movie S1 and Movie S2). These deformed somites then regularly divided into what we call “daughter somites”, and their morphology was consistent with somite divisions observed in N-cadherin and Cadherin-11 knockout mice (Horikawa, Radice, Takeichi, & Chisaka, 1999; Kimura et al., 1995). During daughter-somite formation, an invagination along the mediolateral plane of the deformed somites appeared simultaneously to their separation from the PSM (Fig 1F, Movie S2). It took several hours from the first appearance of this mediolateral invagination for the somite to divide completely. Somite division in stretched embryos appeared unilaterally or bilaterally, and often resulted in daughter somites of different sizes (Movie S2, Fig S4, Fig S5 and Fig S8).

**Fig 1:**
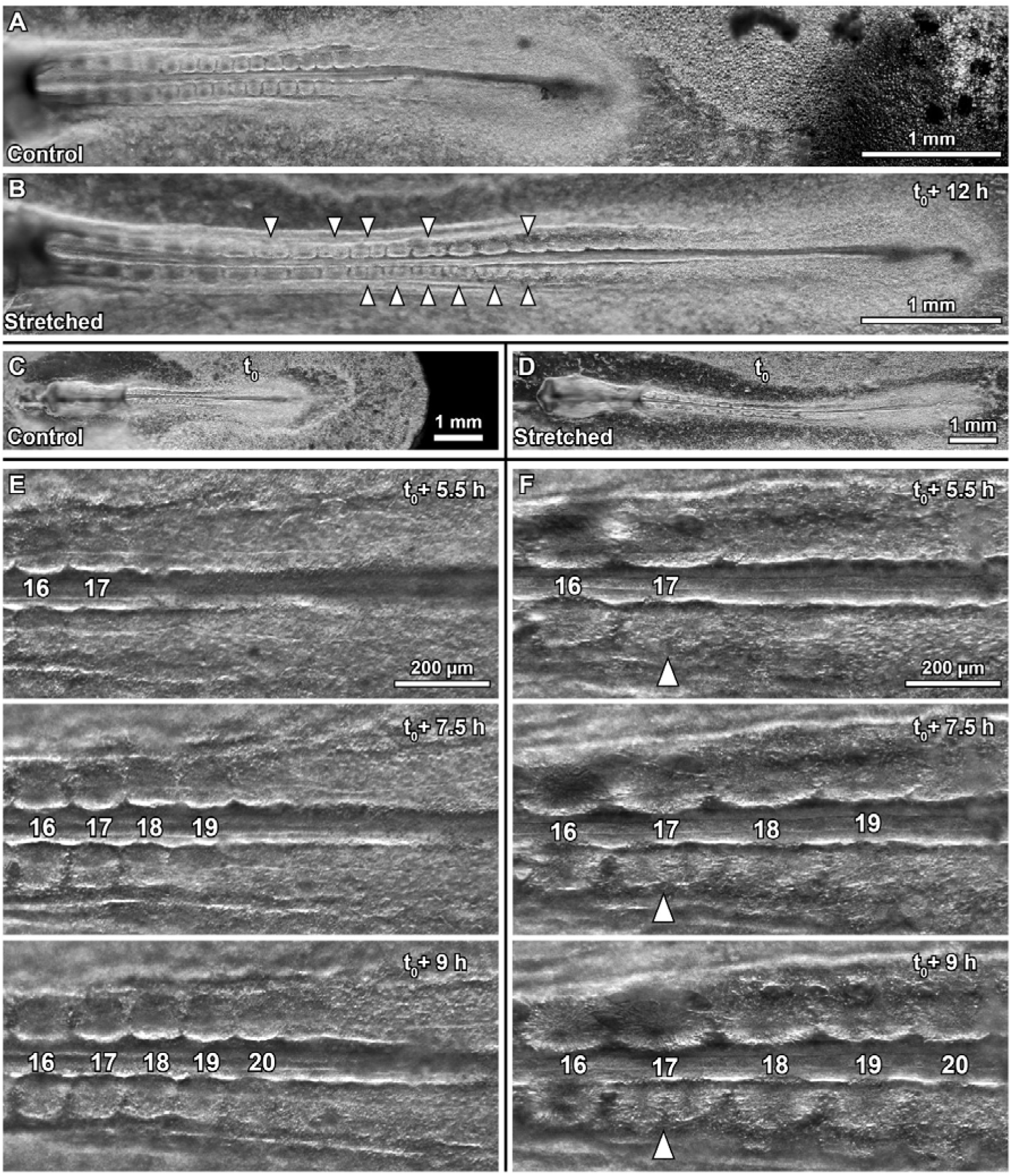
Daughter somite formation in stretched chicken embryos. Dark-field microscopy images of age-matched (**A**) control and (**B**) stretched embryo. Anterior is to the left in all images, arrowheads indicate divided somites, t_0_ = end of stretch protocol, ventral view. Difference in axial length becomes obvious between control embryo (**C**) and stretched embryo (**D**) (t_0_: both embryos are at the 13-somite stage). Selected time-lapse frames of the segmenting PSM in control embryo (**E**) and stretched embryo (**F**). White arrowhead indicates daughter somite formation.

Daughter somites appeared as stable, rounded and clearly separated cellular spheres (Fig 1B). They were morphologically similar to control somites, with epithelial cells organized radially around a somitocoel of mesenchymal cells (compare Fig 2C and G). The daughter somites were enclosed by a fibrous extracellular matrix (ECM) staining positively for fibronectin (Fig S4). While somites in control embryos were rounded (Fig 2A), those in stretched embryos were elongated, while any daughter somites were variable in size (Fig 2B, Fig S4 and Fig S5). Small daughter somites consisting of a few epithelial cells only were also observed (Fig S4). After fixation in transitional stages, the apical actin cortices of somites showed discontinuities along their mediolateral plane, indicating openings of the epithelial sheet under influence of the mechanical deformation (Fig 2D, E and Fig S6). At these locations, mesenchymal somitocoel cells appeared elongated, presumably having undergone mesenchymal-to-epithelial transitions (MET), and were integrated into the existing epithelium (Fig 2E and Fig S6). Upon stretching, there was also strong ectopic expression of *EphA4* in the somitocoels (Fig 2I); this was not present in control embryos (Fig 2H) and indicates stretch-induced MET. During normal somitogenesis, cell-cell signalling between receptor EphA4 in the rostral part of somite S-I and its ligand ephrin B2 in the caudal half of somite S0 has two effects: it induces formation of the somite gap (Watanabe, Sato, Saito, Tadokoro, & Takahashi, 2009), but also establishes epithelialization at somite boundaries by initiating a columnar morphology and cell polarity via apical redistribution of β-catenin (Barrios et al., 2003).

**Fig 2:**
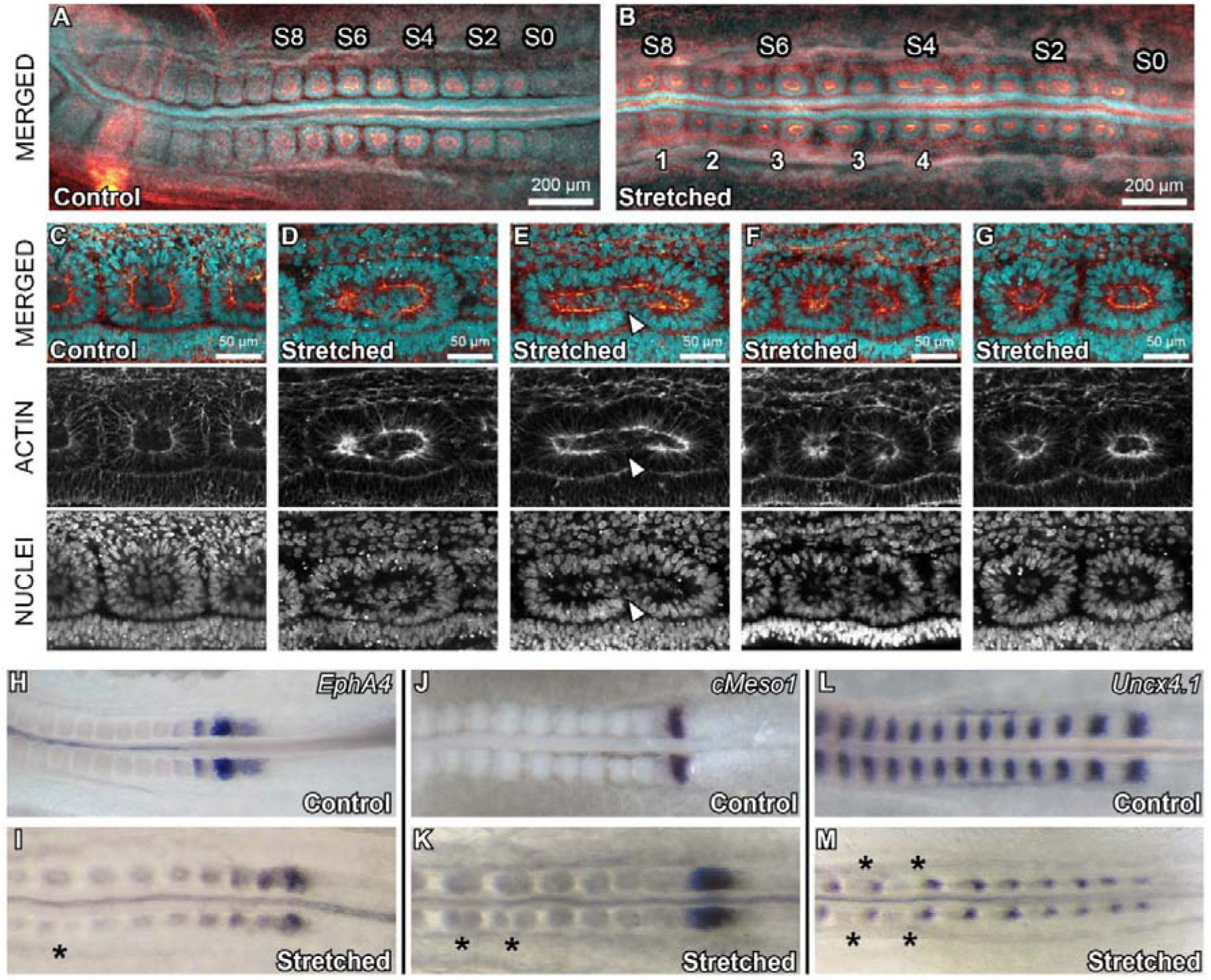
Immunohistochemistry and in situ hybridizations of stretched and control embryos. Control (A) and stretched embryo (B), anterior is left, somite numbers are indicated (Christ & Ordahl, 1995). Scale bars, 200 μm. (B) Daughter somites can form unilaterally (1), equally (4) or unequally sized (3) and subdivide further (2). (C-G) Confocal cross-sections of selected somites of the same control (C) and stretched embryo (D-G). Anterior is left and medial below. Panels are arranged in potential order to illustrate the transition from a mechanically deformed somite (D) to two daughter somites (G). Cells from the somitocoel seem to gain an elongated morphology, possibly before incorporation into the epithelium (arrowhead in E). (H-M) *In situ* hybridizations for *EphA4*, *cMeso1* and *Uncx4.1* show that *EphA4* expression is induced around the somitocoels (I), while no new rostro-caudal polarity is induced in the daughter somites (indicated by *).

However, the expression pattern for *cMeso1*, the key initiator of somite rostro-caudal polarity in the unsegmented PSM in chicken (Morimoto et al., 2007), was not influenced by the stretching (Fig 2K), and the expression pattern of caudal somite marker *Uncx4.1* (Schrägle, Huang, Christ, & Pröls, 2004) showed that daughter somites did not gain a new rostro-caudal genetic identity (Fig 2M). Altogether, our data suggest that mechanical stretching did not affect the unsegmented PSM or the rostro-caudal polarization of the somites. However, mechanical stretching did lead to continuation of EphA4-mediated epithelialization of mesenchymal cells from the somitocoel in somites undergoing daughter-somite formation.

To identify the conditions under which mechanically deformed somites would successfully reorganize into daughter somites, we used the Cellular Potts model (CPM) and implemented a cell-based computer simulation with the open-source package CompuCell3D (Glazier & Graner, 1993; Swat et al., 2012). The somite consisted of a core of non-polarized mesenchymal cells surrounded by a layer of polarized, epithelial cells (Dias et al., 2014). We simulated a somite embedded within an elastic extracellular matrix (ECM; Fig 3A) and mimicked stretching by applying axial tension to the ECM (Fig 3B, Movie S3). To narrow down the possible mechanisms for somite reorganization under mechanical deformation, we tested different rules for cellular behaviour in the deformed somite (Fig S10).

**Fig 3:**
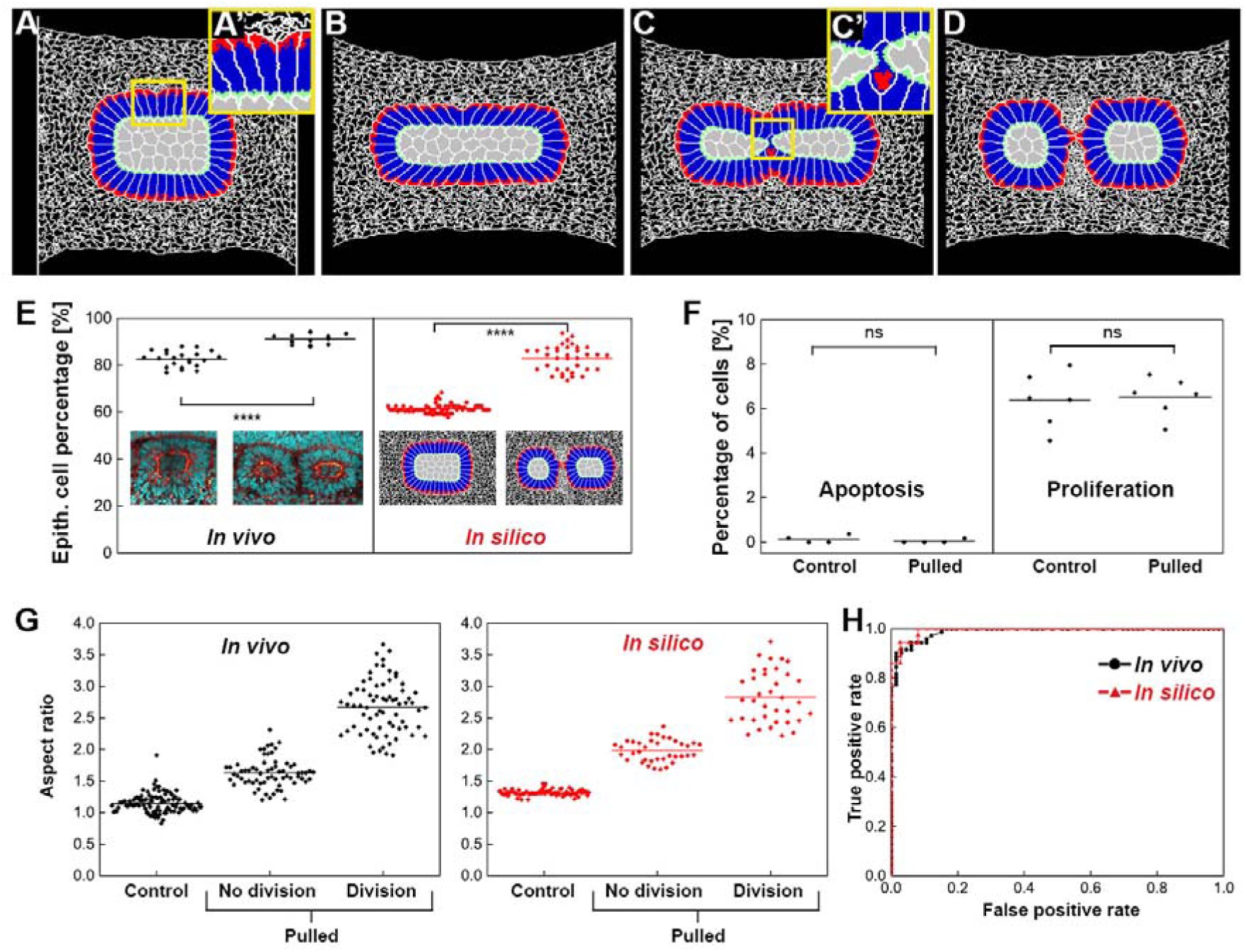
Cellular Potts model of somite remodelling in comparison to daughter somite formation in vivo. (A-D) Daughter somite formation *in silico*, induced by stretching. Mesenchymal cells (grey) and extracellular matrix (white mesh), (A’) epithelial cells consisting of apical (green), lateral (blue) and basal (red) domains. (C’) Somitocoel cells undergoing MET. (E) Percentages of epithelial cells in control and divided somites *in silico* and *in vivo*, average line is shown. Significant increase of epithelial cell fraction *in vivo* (P<0.0001) and *in silico* (P<0.0001). (F) Apoptotic and proliferation rates in somitic mesoderm of control and stretched embryos. Differences non-significant (Mann-Whitney test, apoptosis P=0.43, proliferation P=0.79). (G) Somite aspect ratios *in vivo* and *in silico*. Point clouds and average lines are given. (H) ROC curves for daughter somite formation *in vivo* and *in silico*, in dependence of aspect ratio of deformed somites. AUC is *0.989* (*in vivo*, 95% CI: 0.953-0.999) and 0.993 (*in silico*, 95% CI: 0.938-1.000).

We found that daughter-somite formation *in silico* could successfully be induced only if MET was initiated when mesenchymal core cells came into contact with the basal or lateral membranes of epithelial cells (Fig 3C, C’ and D). To validate this model prediction *in vivo*, we calculated the relative fraction of mesenchymal and epithelial cells in the equatorial cross-section of control somites and divided somites. This showed that the epithelial cell fraction was indeed significantly higher in daughter somites than in controls, and also matched the *in silico* prediction independent of the size of the initial somite (Fig 3E, Fig S11). As the stretching *in vivo* caused no significant changes in the apoptosis and proliferation rates (Fig 3F), we conclude that the increase in the epithelial cell fraction was due to MET.

To determine the relationship between mechanical deformation and somite division, we compared the aspect ratios of stretched dividing somites (prior to division) and stretched non-dividing somites with those of control somites (Fig 3G, Fig S7). These measurements show that somite division is only possible beyond an aspect ratio threshold of about 2 *in vivo* or 2.5 *in silico*. The corresponding receiver operating characteristics (ROC) curves (Goodenough, Rossmann, & Lusted, 1974; Hanley & McNeil, 1982; Lusted, 1971; Metz, 1978) show that the somite aspect ratio is indeed an excellent predictor of somite division *in vivo* and *in silico*, indicating that daughter-somite formation is a highly mechanically-determined process (Fig 3H).

## Discussion

We found that mechanical stretching of live chick embryos can induce a slow reorganization of somites to form two or more well-shaped and stable daughter somites. The complete division of a stretched somite into daughter somites took several hours, indicating that daughter-somite formation is an active process of tissue reorganization, rather than acute mechanical disruption. The resulting formation of new epithelial borders took place in the somitic mesoderm, *i.e.* outside the functional range of the molecular cues described in the clock-and-wavefront mechanism (Hubaud & Pourquié, 2014). Remarkably, the segmentation clock speed (estimated by the average somite formation time) and the genetic segmentation (A-P-polarity of the somites) stay robust during severe deformations imposed on the embryo. As the FGF/WNT gradients are intracellular by mRNA or protein inheritance (Dubrulle & Pourquié, 2004), the cells may retain their local information, despite their different spacing after stretching. Interestingly, our data show that the organization of forming somites is not final after their separation from the anterior tip of the PSM, but that somites can still adapt to their environment.

High-resolution confocal imaging indicates that, during daughter-somite formation, the demand for additional border cells is satisfied by the recruitment of mesenchymal cells from the somitocoel into the existing epithelium. The mechanical deformation creates discontinuities in the apical actin cortices of stretched somites. At the resulting interfaces, mesenchymal cells from the somite core undergo mesenchymal-epithelial transitions (MET) and get integrated into the somitic epithelium. Similarly, it has been shown previously that the development of normal somitic epithelia involves a continuous addition of cells from the somitocoel by accretion and egression (Martins et al., 2009). Our observations indicate that, by adding mesenchymal cells to the epithelium via a similar cellular behaviour during stretching, the epithelium adapts to the changing environment. Under sufficient deformation, this can lead to daughter-somite formation.

We were able to model daughter-somite formation *in silico* only if MET was initiated when mesenchymal core cells came into contact with the basal or lateral membranes of epithelial cells. This suggests a contact-induced mechanism triggering MET of somitocoel cells during daughter-somite formation. *In vivo*, we observed ectopic expression of *EphA4* without c*Meso1* expression in strained somites, although *EphA4* is thought to be downstream of c*Meso1* (Watanabe et al., 2009). This indicates that *EphA4* expression in stretched somites is maintained or reinitiated, suggesting an additional mechanosensitive pathway that leads to *EphA4* upregulation during somite division that is independent from, or redundant to, c*Meso1*. A contact-induced MET mechanism, as suggested here for daughter-somite formation, could underlie general epithelial self-organization during development and homeostasis of epithelia under mechanical stress (Jackson, Kim, Balakrishnan, Stuckenholz, & Davidson, 2017; Martins et al., 2009). This cell behaviour could be mediated via epithelial membrane-based signalling (Baum & Georgiou, 2011; Campbell, Casanova, & Skaer, n.d.), for example on the level of Eph and ephrin binding.

We show that chick somite formation is phenotypically plastic under changing biomechanical conditions. As the geometry of a somite in a stretched embryo predicts with remarkable reliability if it will reorganize into daughter somites or not, daughter-somite formation is highly mechanically determined. This supports the idea that, like temperature, light regime or salinity (Tibblin et al., 2016), mechanical forces can be an additional cue to the induction of different phenotypes of segmental patterning during embryonic development. To further explore the role of mechanical cues in inducing phenotypic plasticity of vertebral numbers, it will be necessary to study the effect of modified mesodermal patterning in stretched embryos on later embryonic development. Though in principle possible (Nagai, Sezaki, Nakamura, & Sheng, 2014), transplanting stretched embryos back into a host egg could prove technically challenging due to the deformation of the embryos. It may therefore be more promising to transplant daughter somites into host chick embryos in the egg and see how skeletal patterning might be influenced.

It has been suggested that the wide range of vertebral numbers between vertebrate species could have evolved for two reasons: (1) The spatial dissociation between axis regionalisation via *Hox* gene expression and segmentation patterning, and (2) the evolvability of the segmentation clock’s period and its relation to the axial growth of the developing embryo (Gomez et al., 2008; Gomez & Pourquie, 2009; Herrgen et al., 2010). However, if the phenotypic plasticity of somite formation shown in our experiments can indeed translate into different vertebral numbers in the later embryo, our findings could indicate an alternative route for vertebrate body plan evolution. Following the ‘side-effect hypothesis’ (G. B. Müller, 2003), natural selection may act on body proportions, leading to changes in the geometry and mechanical loading of the somitic mesoderm. As an initial by-product, the phenotypic plasticity of somites that we show in our experiments then may lead to increasing somite numbers. Later, maintenance of a consistent alteration of the somite number over generations may then become consolidated at the genetic level by natural selection, i.e. via genetic assimilation (Braendle & Flatt, 2006; Fusco & Minelli, 2010), thus providing robustness to the development of the newly acquired body plan. A correlation between axial growth of the embryo and somite size has been shown in mice (Tam, 1981). While daughter-somite formation may be useful as a source of skeletal variation, it remains to be clarified whether and how the developing embryo may cope with the missing A-P polarity of the stretch-induced daughter somites.

Daughter-somite formation could also help to explain an interesting phenomenon. Transitional vertebrae, i.e. vertebrae with morphological characteristic of two adjacent spinal regions (for example lumbar area and sacrum), result from incomplete homeotic shifts of axial identity defined by *Hox* gene expression. Generally, as more than one mutation is needed for complete transformations, incomplete homeotic transformations (and therefore transitional vertebrae) are frequent (Alkema, van der Lugt, Bobeldijk, Berns, & van Lohuizen, 1995; Charité, de Graaff, & Deschamps, 1995; Horan, Wu, Wolgemuth, & Behringer, 1994; Kostic & Capecchi, 1994; Li, Kawasumi, Zhao, Moisyadi, & Yang, 2010; Rancourt, Tsuzuki, & Capecchi, 1995; van den Akker et al., 2001; Varela-Lasheras et al., 2011). However, these transitional vertebrae are statistically underrepresented in several amniote species with variable trunk vertebrae numbers, including *Triturus* newts and lizards ((Kaliontzopoulou, Llorente, & Carretero, 2008; Slijepčević, Galis, Arntzen, & Ivanović, 2015), F. Galis, personal communication, October 2017). This suggests a developmental mechanism which favours complete numbers of trunk vertebrae over transitional vertebrae that might hamper mechanical function (Galis et al., 2014b; Slijepčević et al., 2015). Assuming that additional trunk somites form bilaterally via daughter-somite formation, both newly created somites would have the same axial identity as their mother somite, given that the *Hox* identity is determined mainly before the somite is formed, during the earliest stages of somite formation (Carapuço, Nóvoa, Bobola, & Mallo, 2005; Dubrulle, McGrew, & Pourquié, 2001; Mallo, Wellik, & Deschamps, 2010). This would generate vertebrae of similar regional identity with better mechanical performance than that of potentially disadvantageous transitional vertebrae with a different morphology (Galis et al., 2014a).

Somites’ self-organizational properties provide a promising basis for further exploration into the physical component of somite formation and the causal role of mechanics in body-plan evolution.

## Materials and Methods

### Embryo preparation and culture medium

HH8-9 chicken embryos were explanted using filter paper carriers (Chapman, Collignon, Schoenwolf, & Lumsden, 2001) and cultured *ex ovo* as modified submerged filter paper sandwiches (Chapman et al., 2001; Schmitz et al., 2016), in Pannett-Compton (PC) saline (Pannett & Compton, 1924; Schmitz et al., 2016; Voiculescu, Papanayotou, & Stern, 2008), mixed with freshly harvested thin albumen in a 3:2 ratio. Silicone sheets protected embryos in culture from convection of the medium, thereby avoiding damage. Filter paper carriers were prepared as described recently (Schmitz et al., 2016). Additionally, four holes were cut out from corners of the carriers (Fig S1C) to hook the filter paper sandwiches onto the pins of the motorized arms of the stretching setup (Fig S1A).

### Stretching protocol, axial deformation and somite deformation

Embryos were exposed to a standardized stretching protocol in a custom-made embryo stretcher (Fig S1C). In two consecutive stretching intervals, a slow displacement of the computer-controlled metal arm (see red arrow in Fig S1) extended the filter paper sandwiches by 3.7 to 3.95 mm at a speed of 1.2 μm/s. At this speed, each stretch took 53 to 55 min. We calculated the mechanical strain for the first and the second stretching as relative length change compared to the axial length before stretching. Results are presented in S1 Table. The first stretch lead to 23 ± 3 % strain (average and standard deviation over all 21 embryos presented in S1 Table). The second pull caused 19 ± 3 % strain. For details on the embryo stretching, somite formation time and determination of somite deformation by aspect ratio, see SI Materials and Methods.

### Immunohistochemistry

After the pulling experiments, the embryos and age-matched controls were fixed in 4% paraformaldehyde overnight in PBS at 4°C. Permeabilization in PBST + 0.15% Triton-X-100 lasted for 1.5 hours. Blocking was performed for 2 hours in PBST + 2% BSA + 5% normal goat serum. The following antibody was used: fibronectin mouse-anti-chicken (B3/D6-s, Hybridoma bank). The antibody was diluted in PBST with 1% BSA. Embryos were incubated in primary antibody solution for 24h at 4°C, followed by extensive washing in PBS and incubation with appropriate Alexa Fluor-conjugated secondary antibody (1:500, Molecular Probes). Embryos were stained for F-actin using Alexa Fluor 546 Phalloidin (1:200, Molecular Probes) and for nucleic DNA using DAPI (1 μg/ml). Cell proliferation and apoptosis staining was performed using following antibodies: rabbit polyclonal anti-cleaved caspase-3 (1:200, Cell Signaling) and rabbit polyclonal anti-phosphohistone-H3 (1:400, Cell Signaling) with the appropriate Alexa Fluor-conjugated secondary antibodies (1:500, Invitrogen) and DAPI for nucleic DNA (1 μg/ml). For details on mesenchymal and epithelial cell counts, see SI Materials and Methods.

### In situ hybridizations

In situ hybridizations were performed by standard procedures. Embryos were fixated in freshly prepared 4 % PFA in PBS. The embryos were pre-treated with proteinase-K in PBST at 37°C with agitation for 3 minutes. During staining, embryos were incubated in NTMT containing 4.5 μl NBT (75mg/ml in 70% DMF) and 3.5 μl BCIP (50mg/ml in 100% DMF) per 1.5 ml. Pulled embryos and age-matched controls were stained in the same wells for the same time, as much as possible. After the staining had been stopped, the embryos were photographed in glycerol 80% in H_2_O with a Leica DFC320 camera on a Leica MZ75 microscope.

### Cellular Potts model of somite division

To understand the influence of mechanical stretching on somites and the somite division observed *in vivo*, we constructed a two-dimensional mathematical model based on the Cellular Potts Model (Glazier & Graner, 1993; Graner & Glazier, 1992), representing a cross-section through a three-dimensional somitic tissue. For the modelling details, see SI Materials and Methods.

## Acknowledgements

We are grateful to Julio Belmonte for his advice on the simulations at the early stages of model development. We thank Stuart A. Newman and Frietson Galis for valuable comments during the final preparation of the manuscript.

## Author Contributions

B.N. and M.S. performed the experiments and analysed the data. H.T. and R.M. performed the simulations. T.S. and R.M. supervised the project. All authors wrote the manuscript.

## Supporting Information

### SI Materials and Methods

#### Egg handling

Fertilized chicken eggs, white-leghorn, *Gallus gallus domesticus* (Linnaeus, 1758), were obtained from Drost B.V. (Loosdrecht, The Netherlands), incubated at 37,5°C in a moist atmosphere, and automatically turned every hour. After incubation for approx. 33 h, HH8-9 embryos were explanted using filter paper carriers (Chapman et al., 2001) and cultured *ex ovo* as modified submerged filter paper sandwiches (Chapman et al., 2001; Schmitz et al., 2016)

#### Culture medium

Embryo culture medium consisted of Pannett-Compton (PC) saline (Pannett & Compton, 1924; Schmitz et al., 2016; Voiculescu et al., 2008), mixed with freshly harvested thin albumen in a 3:2 ratio. PC stock solutions can be stored at 4 °C for several months, but PC saline (mixture of stock solutions and MilliQ-water) should be prepared freshly every week and stored at 4°C between experiments. Addition of Penicillin/Streptomycin (10000 U/ml) in 100x dilution prevents occasionally appearing bacterial infections.

#### Silicone sheets

Silicone sheets protected embryos in culture from convection of the medium, thereby avoiding additional damage. Silicone sheets (ca. 350 μm in thickness) were made using a Sylgard^®^ 184 Silicone Elastomer Kit as follows: A 15-cm plastic petri dish was placed on a scale and 6.165 g (5.554 mL) of base solution were pipetted into its center using a plastic transfer pipette (cut off tip). Then 0.206 g (0.2 mL) of curing agent were added using a glass pipette. Base and curing agent were mixed slowly, using a wooden spatula and spread out over the bottom of the petri dish. The petri dish was placed into a vacuum chamber for 2 hrs to remove air bubbles and let silicone solution spread out equally. Exposure to 80 °C for ca. 2 hrs let silicone polymerize and cure. Afterwards, silicone was let to cool to room temperature for about 5 hrs or overnight. Tweezers were used to free the borders of the silicone sheet from the walls of the petri dish and peel the sheet from the culture dish (wear gloves). Silicone was stored between sheets of a plastic document sleeve to prevent accumulation of dust. For preparing silicone sheets fitting in the setup, the plastic sleeve was removed from one side of the silicone sheet and the plastic stencil (Fig S1D) placed on it. A razor blade was used to cut around the outline of the stencil and the ten holes indicated by the stencil were cut out using a hole puncher. After removing the other plastic sleeve layer, silicone sheets were stored in a closed 10 cm petri dish.

#### Filter paper carriers

Filter paper carriers were prepared as described recently (Schmitz et al., 2016). Additionally, four holes were cut out from corners of the carriers (Fig S1C) to hook the filter paper sandwiches onto the pins of the motorized arms of the stretching setup (Fig S1A).

#### Experimental setup - Embryo stretcher

Embryos were cultured and mechanically manipulated on a custom-made embryo stretcher (Fig S1C). This setup allows to culture up to three embryos simultaneously in a variant of the recently described “submerged filter paper sandwich” (Schmitz et al., 2016). The setup consisted of a temperature-controlled medium container/ beaker surrounded by a metal frame. This frame carried two motorized translation stages mounted on opposing sides of the frame. Both stages could be operated with a manual controller. For one of the stages the control could be switched to a custom-made LabVIEW routine. This routine allowed to define different automated pulling profiles for overnight experiments (see section ‘Stretching protocol’). Each stage carried a metal arm reaching into the medium container. Each metal arm ended with a horizontal platform equipped with three arrays of four smooth pins surrounding a threaded pin. The smooth pins functioned as hooks for the filter paper sandwiches and as a guide for metal washers clamping the filter paper sandwiches to the metal arms. Embryos were cultured fully submerged in the culture medium described above and the setup was prepared for an experiment as follows: The temperature-controlled beaker was placed in the center of the motorized x-y-stage of the upright zoom microscope. Then the frame carrying the two motorized translational stages was placed around the temperature-controlled beaker and fixed it with two lateral screws (Fig S1B). The two metal arms were attached to the motorized translational stages and fixed with the screws from top. Using the manual control, the position of the metal arms was adjusted so that the gap between them was 12 mm, by bringing the left translational stage furthest to the right and the right translational stage furthest to the left. Then the temperature controlled beaker was filled with 200 mL of clean culture medium. Any dirt or bubbles were removed from the culture medium using a plastic transfer pipette. If necessary, the culture medium level was adjusted to make sure that the pins on the metal arms were just submerged. The temperature of the temperature-controlled beaker was set to 40°C.Then the silicone sheets were placed into the setup by using the blunt end of tweezers to hook the silicone sheets into the pins of the metal arms. Air bubbles were removed with a plastic transfer pipette. Then a chick embryo was explanted into a filter paper sandwich (Schmitz et al., 2016) (Fig S1A), immediately submerged into the culture medium and hooked it into the innermost pins of both metal arms by pushing it down with the tweezers (Fig S1A The metal washers (Fig S1A and Fig S1E for dimensions) were placed over the pins of both metal arms and gently fixed by nuts using an Inbus^®^ key (Fig S1A). After successfully clamping three filter paper sandwiches into the setup, each filter paper sandwich was cut perpendicularly to the embryonic axis, about 1 mm posteriorly of the widest point of the elliptical aperture (dashed red line in Fig S1A-9). Iris scissors were used for these cuts. Then the surface of the culture medium was covered with 50 mL of light mineral oil using a plastic transfer pipette (Fig S1A-10). Note: After placing the mineral oil, the experiment is set and embryos cannot be replaced anymore without cleaning the whole setup. 25 mL of culture medium were removed from below the oil layer by pinching a plastic transfer pipette through the oil layer into the culture medium and the time-lapse acquisition was set up as described before (Schmitz et al., 2016).

#### Stretching protocol

Embryos were exposed to a standardized stretching protocol which, in preliminary experiments, had proved to cause a strong deformation of the embryos’ paraxial mesoderm without impairing the progression of development. The protocol started with a waiting period of 3.5 hrs after covering the culture medium with light mineral oil. This allowed damage, caused during preparation of the filter paper sandwich and perpendicular cutting, to heal. Then, two consecutive stretching intervals followed, separated by a pause of two hours to allow recovery of damaged tissue. Each stretching interval consisted of a slow displacement of the computer controlled metal arm (see red arrow in Fig S1) by 3.7 to 3.95 mm at a speed of 1.2 μm/s. At this speed, each stretch took 53 to 55 min. Fig S2 depicts the user interface of the LabVIEW routine, prepared for an overnight experiment. Travel direction (left or right), speed (to be chosen from a 16-bit range defined by the microcontroller, where 40 equals 1.2 μm/s) and time interval (in seconds) could be chosen independently for each experimental block.

#### Macroscopic axial deformation

The macroscopic axial deformation of the stretched embryos was determined by measuring their length from tip of the head to the posterior end of the sinus rhomboidalis, using the segmented line tool in ImageJ (white line in Fig S3A, B, E, F). We assumed that, during the application of the stretching, the natural morphogenetic changes of the embryos are negligible and length changes result from the external stretching only. We calculated the mechanical strain for the first and the second stretching as relative length change compared to the axial length before stretching. Results are presented in S1 Table. The first stretch lead to 23 ± 3 % strain (average and standard deviation over all 21 embryos presented in S1 Table). The second pull caused 19 ± 3 % strain.

#### Somite formation time

The separation of a newly forming somite from the anterior tip of the PSM and its epithelialization is a continuous process involving complex cellular rearrangements lasting longer than the 90 min usually stated to be characteristic for chick embryos (Martins et al., 2009). Nevertheless, we tried to assess the influence of mechanical deformation on somitogenesis by determining an average somite formation rate for stretched and control embryos using our dark field microscopic time-lapse movies and based our calculations on the physical separation of somites from the PSM. We counted the number of somites in stretched embryos at the end of the second pull and at the end of the experiment. If a somite had not completely separated from the PSM at the end of the second pull, the counting was started after formation of the following somite. From the total number of somites formed after the application of the second pull and the corresponding time interval we calculated the somite formation rate. The somite formation rate for control embryos was determined accordingly from the beginning of the culturing in the submerged filter paper sandwich. We did not observe any difference in somite formation rate between stretched embryos with or without daughter formation or in comparison to non-stretched control embryos (S2 Table).

#### Proliferation rate and apoptotic rate

Apoptotic (Cas3 staining, control n=4, pulled n=4 embryos) and proliferating (pHH3 staining, control n=6, pulled n=6 embryos) cells in somitic mesoderm lanes (somite S1 to S5) were counted in high-resolution confocal micrographs acquired on a Leica SP8 confocal microscope. At least 500 cells were counted per embryo. Apoptotic rate and proliferation rate were calculated as follows: (apoptotic/proliferation) rate (%) = number of positive staining cells/number of total cells×100. Statistical analysis was performed using GraphPad Prism software. Mann–Whitney unpaired non-parametric two-tail testing was applied to determine the P-values for the apoptotic and proliferation rates shown in Fig 3F. Control and pulled somitic mesoderm showed no significant differences in apoptotic (P=0.4286) and proliferation (P=0.7879) rates.

#### Epithelial cell percentages

The percentage of epithelial cells in the equatorial z-plane of 13 daughter somite pairs, originating from the same mother somite, and 22 control somites was determined (*in vivo*). To that end, high-resolution confocal micrographs of embryos stained with DAPI for nucleic DNA were acquired on a Leica SP8 confocal microscope. Then somites were counted for (mesenchymal) core cells and epithelial cells to calculate epithelial cell percentages. *In silico*, cell percentages in 12 daughter somite pairs and 12 control somites were counted accordingly. Statistical analysis was performed using GraphPad Prism software. Unpaired parametric two tailed t-tests (with Welch’s correction for unequal variance) were applied to determine P-values for the epithelial percentages shown in the graph in Fig 3E. The percentages of epithelial cells change significantly *in vivo* (P<0,0001) and *in silico* (P<0,0001).

#### Aspect ratio determination and ROC curve

The geometry of somites *in vivo* in controls and stretched embryos was described by measuring their length in rostro-caudal (x) and their width in medio-lateral (y) direction using the “Measure”-tool in ImageJ. Subsequently, the corresponding aspect ratio AR (AR = x/y) was calculated. Somites forming in controls and in stretched embryos after the second pull were measured upon their separation from the anterior tip of the PSM (Fig S7A-B). Somites that had been formed before were measured at the end of the second pull (Fig S7C). The aspect ratio of somites *in silico* was determined before (Fig S7D) and after the application of the pull (Fig S7E) accordingly (for strain regimes *in silico* see below). The corresponding receiver operating characteristics (ROC) curves (Goodenough et al., 1974; Hanley & McNeil, 1982; Lusted, 1971; Metz, 1978) (Fig 3H) were generated by performing a binary logistic regression using the Data Analysis Tool of the Real Statistics Excel plugin Realstats (available at http://www.real-statistics.com). We analyzed how well the aspect ratio of stretched somites *in vivo* and *in silico* can predict the binary outcome of whether a somite will undergo division or not. This is measured by the area under the curve (AUC) in the ROC diagram. The AUC can very between 0.5 (stochastic relation) and 1 (fully determined). 95% Confidence intervals for AUC values were calculated using the ‘ROC curve analysis’ tool of MedCalc software (available at https://www.medcalc.org/index.php).

#### Cellular Potts model of somite division

To understand the influence of mechanical stretching on somites and the somite division observed *in vivo*, we constructed a two-dimensional mathematical model based on the Cellular Potts Model (Glazier & Graner, 1993; Graner & Glazier, 1992), representing a cross-section through a three-dimensional somitic tissue.

We made use of a compartmentalized Cellular Potts model (CPM), also known as Glazier-Graner-Hogeweg model. The model was implemented using CompuCell3D, an open source modeling package based on the CPM (Swat et al., 2012). CompuCell3D’s ‘compartmentalized cell’ module implements the compartmental CPM that represents biological cells as a collection of sub-cellular domains, each identified by a unique cluster index. This module allows specification of separate contact energies between the domains of a single cell (internal parameters) as well as specification of external contact energy parameters between cells of the same type and of different types. For more information on CPM modeling or compartmentalized cell module, please refer to Swat *et al.* (Swat et al., 2012).

Briefly, the compartmentalized CPM projects biological cells on a (usually regular, rectangular) lattice as domains of (usually) connected lattice sites. Each lattice site, 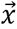, is associated with a domain index 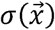 that identifies a biological cell, a cellular compartment, or a volume element of extracellular material. Cell identification number *σ* = 0 usually represents a generic ‘medium’. Each domain *σ* has a type label 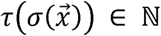 to represent the generic ‘type’ (subcellular domain, ECM, and so forth) and an additional label 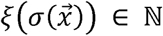 that bundles compartments to a biological cell or connected extracellular material. Although each individual object (subcellular compartment, ECM medium etc.) has its own unique domain index *σ*, many objects may be associated with the same type label *τ*.

The evolution of our CPM is governed by a force-balance, represented by a Hamiltonian, *H*,

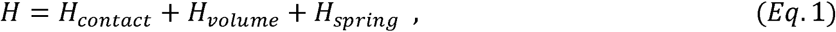

which describes the dynamics of cells (e.g. cell behaviors, properties and interactions). The Hamiltonian is minimized using a Metropolis algorithm that mimics microscopic membrane and material fluctuations, such that both the equilibrium and the transient towards the equilibrium can be physically and biologically interpreted (Newman & Barkema, 1999). *H*_*contact*_ represents the cell adhesion where cell-cell and cell-medium interactions take place through contact energies. The size of the interface between two cells defines the contact energy and is given by:

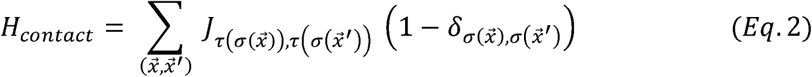

Here, 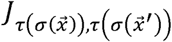 is the bonding energy between two neighboring cell types 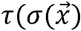 and 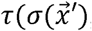, and 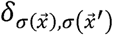 is the Kronecker delta term in which adhesion is restricted to the cell membranes by eliminating the contributions from the neighboring lattice sites belonging to the same cell. If 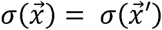, the delta function returns a value of 1 and 0 otherwise. The term *H*_*volume*_ in the Hamiltonian specified in Eq. 1 is given by:

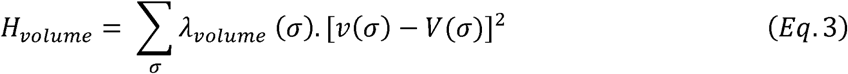

and constrains the cell volume, *v*(*σ*), close to a resting volume *V*(*σ*). The Lagrange multiplier *λ*_*volume*_ represents cell elasticity - higher values of *λ*_*volume*_ reduce fluctuations of cell’s volume from its target volume.

Compartments of cells and subunits of the extracellular matrix can be mechanically coupled by connecting their centers of mass using springs. Each spring contributes an additional energy bias *H*_*spring*_ to the Hamiltonian in *Eq. S2*.

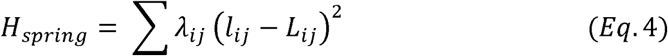

where *l*_*ij*_ is the absolute distance between the center of masses of cells *i* and *j* and *L*_*ij*_ is the specified target spring length. *λ*_*ij*_ is an elasticity parameter. Springs rupture if they exceed a threshold length; new springs are formed if cells move within a threshold distance. In our simulations, we have many cell types (epithelial internal compartments, extracellular matrix (ECM), and epithelial cell (apical)) which are connected using springs. Details on these cell-specific spring lengths and elasticity are explained in the next sections of the supplementary material.

The CPM is updated using a Metropolis algorithm, which mimics the extension and retraction of pseudopods of the biological cells, and fluctuations of the extracellular matrix materials. The algorithm iteratively selects a lattice site 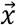 and attempts to copy its cell index 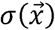 into a randomly chosen adjacent lattice site 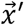. This is called a copy attempt. The probability of accepting or rejecting the attempted copy update is based on the energy minimization criteria and follows Boltzmann probability,

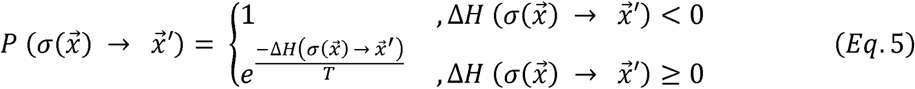

where 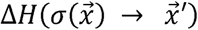 represents the change in the Hamiltonian due to the copy attempt. *Eq. 5* shows that if the attempted copy update will reduce the energy, i.e. 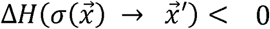, the update is accepted with a probability of 1. However, if the energy increases due to the copy-attempt, the system follows Boltzmann probability to accept or reject a copy-attempt. The parameter *T* is the cellular temperature, representing the amplitude of active cell membrane fluctuations or fluctuations of the extracellular materials.

The model included the following assumptions: (i) the tissue surrounding the somite can be approximated as elastic, and was modeled as a non-specified extracellular matrix (ECM); (ii) the somite consists of polarized epithelial cells forming the outer layer (Dias et al., 2014), while the somite core (somitocoel) consists of unpolarized mesenchymal cells. The mesenchymal cells in the core of the somite were represented by single-compartment, non-coupled and non-polarized cells. Following Dias *et al.* (Dias et al., 2014) the epithelial cells in our model consisted of three domains, called ‘apical’, and ‘lateral’ and ‘basal’ (Fig S9). The three compartments were initially distributed at random inside an epithelial cell and after a brief relaxation period of 1500 MCS (epithelial polarization time), these compartments were connected internally to one another using linear elastic springs (Eq. 4). To achieve epithelial elongation (Fig S9), target lengths of all internal springs (*L*_*apical-lateral*_, *L*_*apical-basal*_, *L*_*lateral-basal*_) were individually incremented by 1 every 20^th^ MCS within the elongation time frame (1500 MCS), until every spring reached its final specified target length (*L*_*apical-lateral*_ = 15, *L*_*apical-basal*_ = 30, *L*_*lateral-basal*_ = 20). The ECM, with its main functional component fibronectin *in vivo* (“P.Rifes, thesis: Fibronectin cues during somite formation, Universidade de Lisboa (2013),” n.d.), was modeled as a network of cells. These ECM cells were also connected to each other using elastic springs of target length *L*_*ecm-ecm*_ = 10 with elastic stiffness *λ*_*ecm-ecm*_ = 200. This was necessary to obtain a desired stiffness of the ECM in order to keep it intact during and after stretching.

We first attempted to construct a well-organized, initial epithelial structure as a starting point for the stretching model. Contact energies between domains as well as contact energies with other cell types and the ECM were set according to S3 Table. In absence of quantitative values for the adhesion strengths and interfacial tensions between the cells, we estimated parameter values for which a stable epithelial monolayer is maintained in our simulations, followed by parameter sensitivity studies. We assumed the apical domains of adjacent epithelial cohered strongly, following Dias *et al.*’s (Dias et al., 2014) assumption mimicking the distribution of N-Cadherin *in vivo* (Chal, Guillot, & Pourquié, 2016). The lateral domains of epithelial cells (between the apical and basal domains) adhere strongly to each other, similar to Cadherin mediated cohesion *in vivo* (Horikawa et al., 1999; Kimura et al., 1995). To represent the apical actin ring, each center of mass of an apical unit was connected to the center of mass of neighboring apical domains on either side (left and right) using elastic springs of a resting length of *L*_*ij*_ = 4 with elastic stiffness *λ*_*apical-apical*_ = 100.

The monolayer of epithelial cells was constructed by initializing the simulation with a collection of mesenchymal cells surrounded by an elastic ECM. We selected a mesenchymal cell at the boundary with the surrounding ECM and made it epithelial. This first epithelialized cell induced MET in neighboring cells based on basolateral contact (Baum & Georgiou, 2011; Campbell et al., n.d.), which finally led to a fully epithelialized somite-like structure (initial phase of Movie S3).

After a stable, somite-like, epithelial structure had formed, we gradually strained the extracellular matrix in our simulations, in order to mimic the experimental setup. To this end, we connected two ‘walls’ constructed out of immobile cells to the left and right-hand ends of the ECM using stiff elastic springs and slowly moved the walls apart by 1 pixel every 50 MCS, using a technique introduced previously for simulating the compression of tissue spheroids (Marmottant et al., 2009) and the application of stents in arteries (Tahir, Niculescu, Bona-Casas, Merks, & Hoekstra, 2015)

The stretching rate was sufficiently slow (walls moved outward by 1 pixel every 50 MCS), such that it did not damage the ECM cells (Phase 2 in Movie S3). We tested three MET scenarios for their ability to allow the stretched somite to divide into epithelialized daughter somites. We first assumed that somites can divide by reorganization of existing epithelial cells and allowed no MET after the stretching. We could successfully deform the somite by stretching, but neither small nor large strains induced somite division (Fig S10, top row). This suggested the necessity of an MET mechanism to enable somite division. In the second scenario, a mesenchymal cell underwent MET after contact with the surrounding ECM for a certain time (600 MCS). We did not observe somite division for different levels of deformation, probably because intercellular connections between epithelial cells did not soften completely upon stretching, thereby avoiding sufficient contact between mesenchymal somitocoel cells and the surrounding ECM to induce additional MET (Fig S10, middle row). In the third scenario, the MET occurred after sufficiently long contact of mesenchymal cells to the basal or lateral membrane of epithelial cells (600 MCS). Upon stretching, several springs between neighboring apical compartments released and mesenchymal cells from the core became exposed to the lateral or basal membranes of epithelial cells leading to additional MET. These additional epithelial cells disturbed the equilibrium and could not get incorporated into the original epithelial ring. So, the epithelium started to reorganize and divide into daughter somites (Fig S10, bottom row and Phase 3 in Movie S3).

Based on our observations that daughter somites are separated and presumably stabilized by a newly forming fibronectin matrix (Fig S4), we also implemented a similar rule for ECM production by the basal units of epithelial cells. If the basal domain of an epithelial cells is not attached to a specified amount of ECM (given by threshold value) for a certain duration, it produces an additional ECM cell. This production continues until the threshold value is reached again. Such production of the fibronectin allows the dividing somites to separate from each other permanently. The parameters used in the simulations are shown in the S3 Table. For a systematical analysis of how well the geometry of stretched somite predicts division we applied lateral wall displacements of 30 to 110 pixels (15 to 55 pixels per side), resulting into aspect ratio values similar to stretched somites *in vivo* (Fig 3G).

The *in silico* somite model can be parameterized to the experimentally observed ratio of mesenchymal and epithelial cells. Fig S11 shows that, following the MET based on the basolateral contact rule, initial epithelialization of the somite and division of the somites after stretching can be achieved with a large core and small core somite.

In order to further validate the *in silico* model, we also tested the influence of decreased cohesion between lateral domains of epithelial cells in epithelializing, non-stretched somites. Similar to results in *N-Cadherin/cad11* double-homozygous mouse mutants (Horikawa et al., 1999), we observed subdivisions into small cell clusters of epithelioid morphology (Fig S12).

---

**Fig S1:**
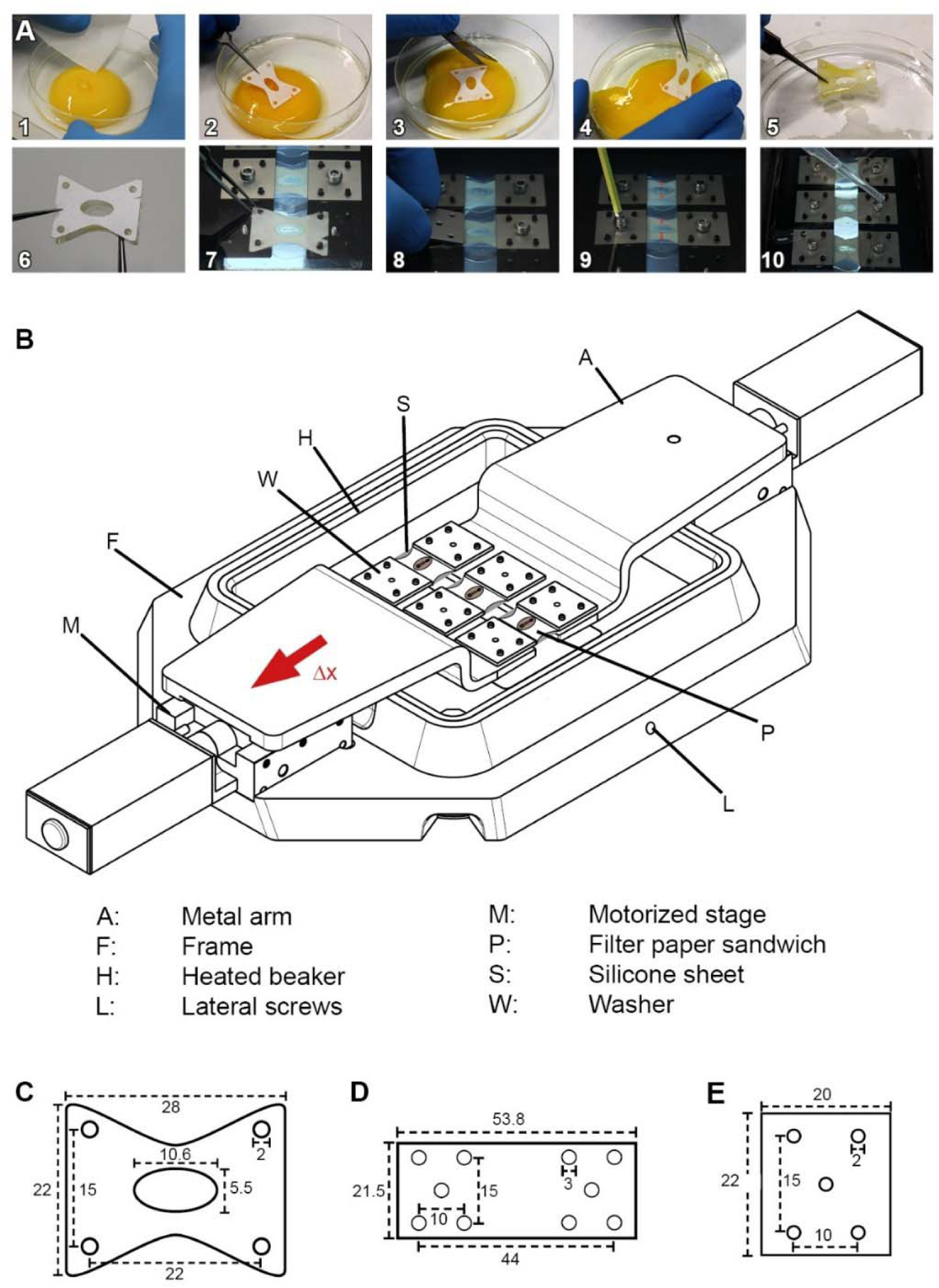
Chick embryo stretching *ex ovo*, experimental setup. (A) Explantation procedure (Schmitz et al., 2016). (1) An egg is cracked into petri-dish and thick albumen removed from top of the yolk. (2) A filter paper carrier is placed on top of the yolk, surrounding the blastoderm. (3) The filter paper carrier is cut loose from the surrounding vitelline membrane and (4) removed from top of the yolk. (5) Remaining yolk is carefully washed away in a saline bath. (6) The embryo is sandwiched with a second filter paper carrier. (7) The filter paper sandwich is submerged into the medium and hooked into the pins of the motorized arms. A thin sheet of PDMS protects the embryo from convection of the medium. (8) Washer plates clamp the filter paper sandwich to the metal arms and (9) are carefully pressed down by nuts. Filter paper sandwiches are cut along dashed red lines for later stretching of embryos. (10) The medium is covered with a layer of light mineral oil. (B) Schematic view of the embryo stretcher. The frame carries the motorized stages and keeps the temperature-controlled medium container in position. The whole setup is placed on a motorized x-y-stage, embryos are imaged from above and illuminated from below through the glass bottom of the medium container. (C) Filter paper carrier dimensions (in mm). (D) Dimensions of stencil for PDMS sheets (in mm). (E) Dimensions of metal washers used to clamp the filter paper (in mm).

**Fig S2:**
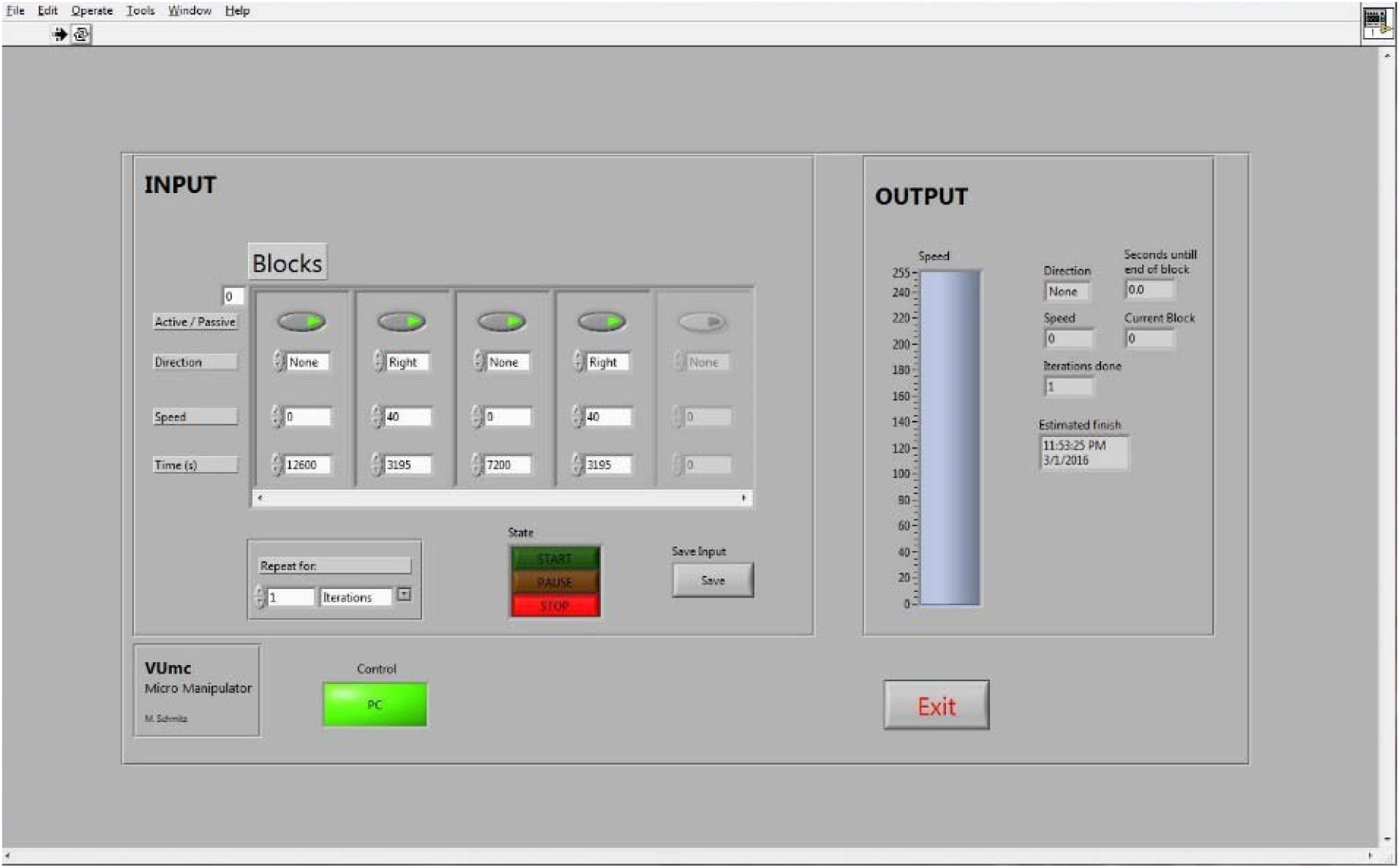
User interface of the custom-made LabVIEW routine used to control one motorized translation stage of the embryo stretcher. Four experimental blocks are activated, defining the resting time after covering the culture medium with light mineral oil (12600 s/3.5 hrs), the first pulling block (3195 s), a resting time in between pulls (7200 s/2 hrs) and the second pulling block (3195 s).

**Fig S3:**
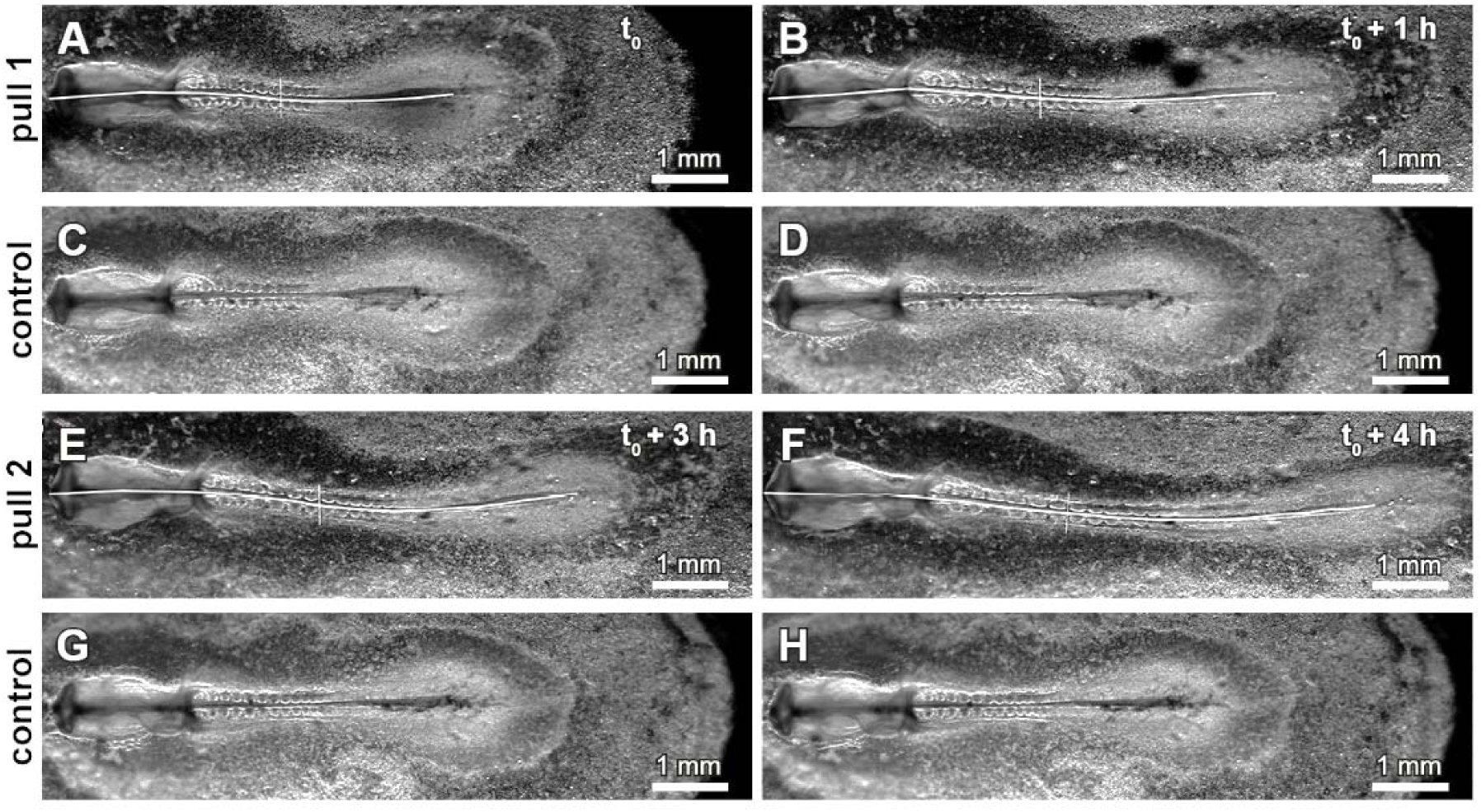
Macroscopic deformation of pulled embryos. Before application of the first pull, experimental embryo (A) and control (C) are similar in axial length. The first pull increases the axial length of the pulled embryo considerably compared to the control (compare B and D). This length difference remains over the waiting period between first and second pull (compare B and D and E and G). The second pull increases the length difference between pulled embryo and control even further (compare F and H).

**Fig S4:**
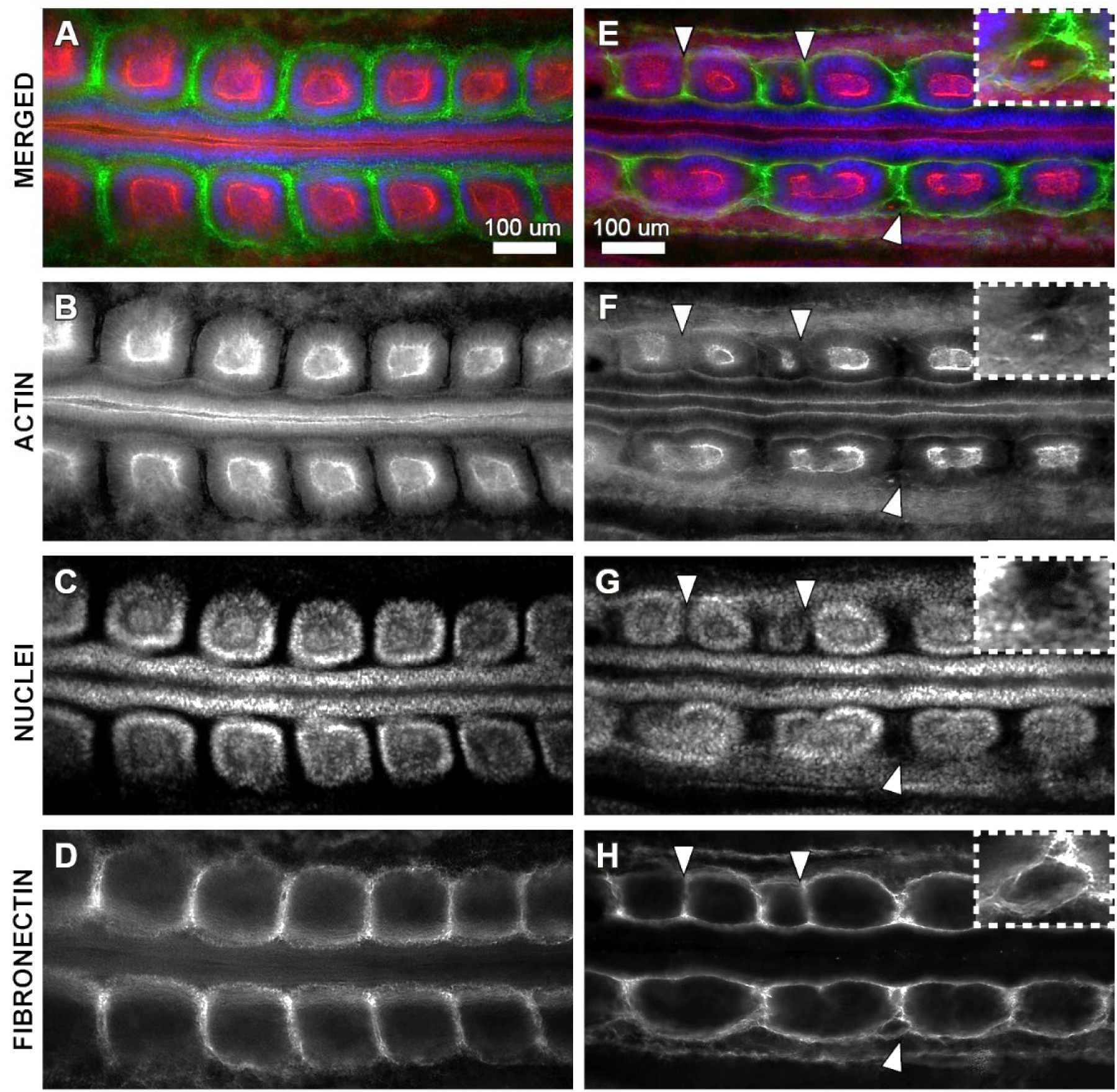
Fibronectin distribution around daughter somites. Widefield fluorescent micrographs of control embryo (A-D) and stretched embryo (E-H) stained for actin (red), DNA in cell nuclei (blue) and extracellular matrix component fibronectin (green). Ventral view, anterior is to the left. Daughter somites are surrounded and separated from each other by a newly formed fibronectin matrix (white arrowheads in E and H) and can be extremely small (inset E to H) and lacking a mesenchymal core.

**Fig S5:**
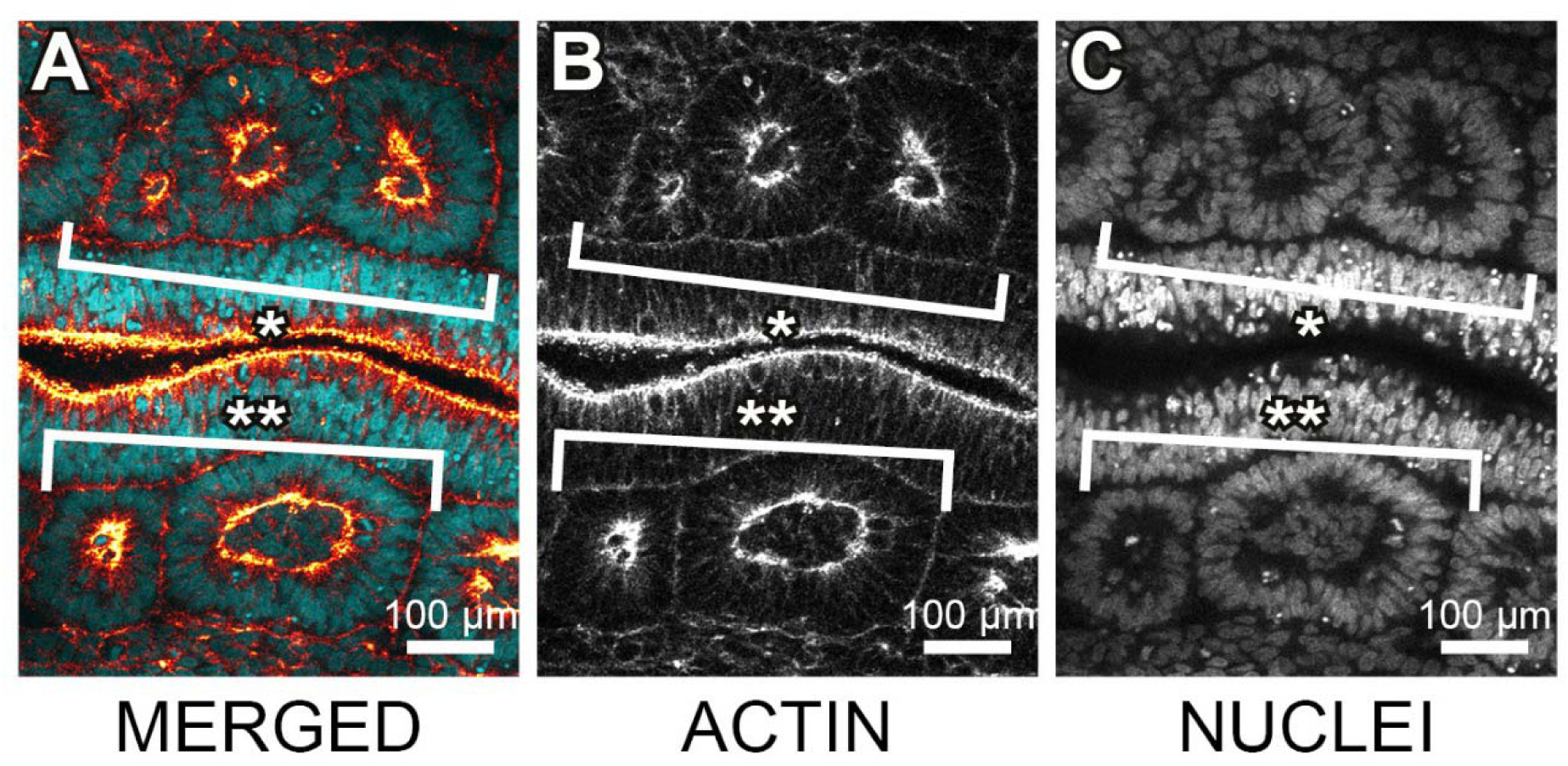
Versatile daughter somite morphologies. Confocal micrograph of daughter somites on both sides of the midline in stretched and fixated chick embryo stained for actin (red) and DNA in cell nuclei (blue). Anterior is to the left, ventral view. Daughter somites do not only result from splitting into anterior and posterior compartment of original somite, but can also reorganize into three daughter somites (*) or two unequally sized daughter somites (**).

**Fig S6:**
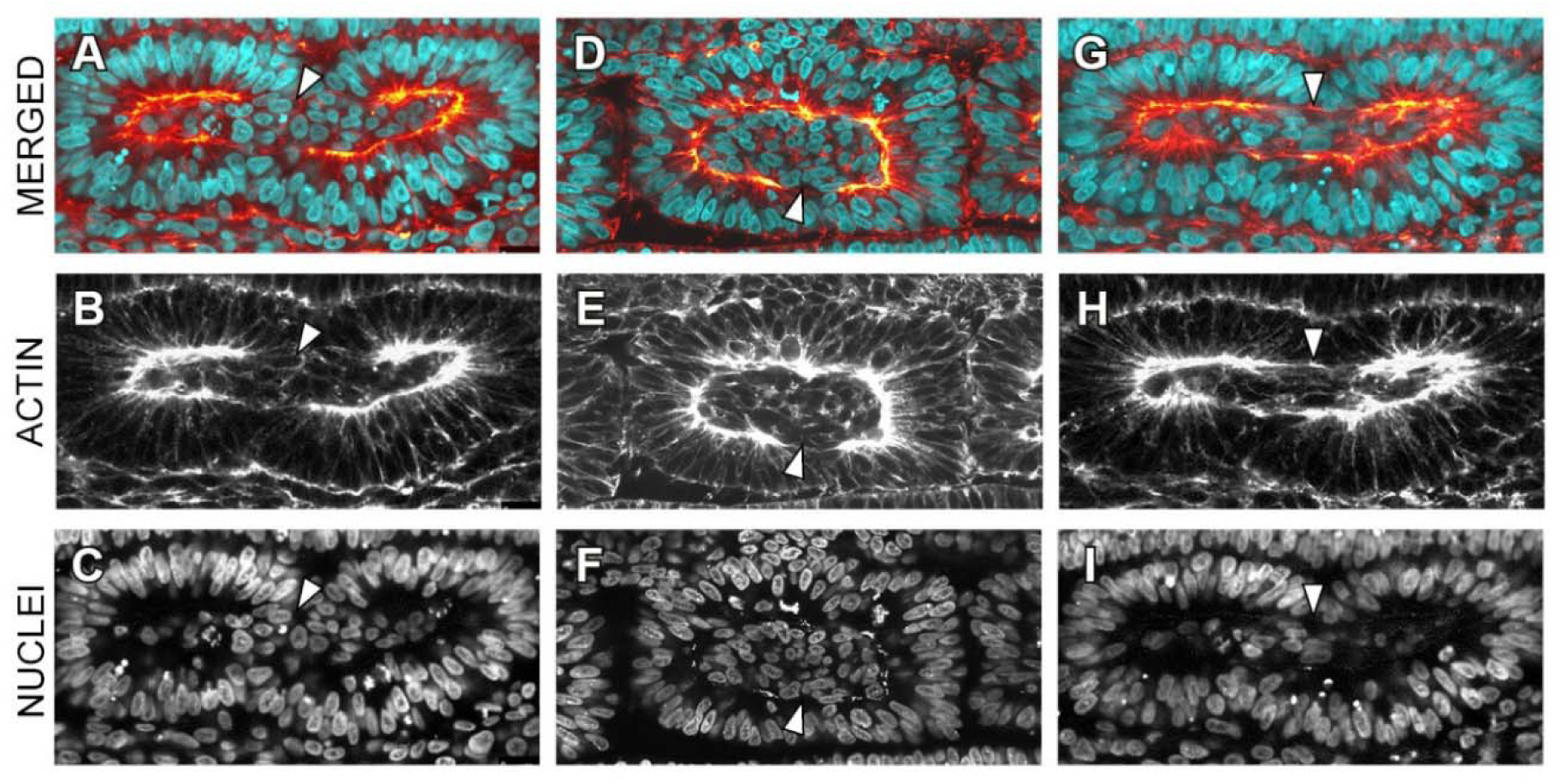
Fracture of epithelial sheet and potential MET of mesenchymal somitocoel cells. Confocal micrograph of somites, fixated during daughter somite formation in stretched chick embryos, stained for actin (orange) and DNA in cell nuclei (cyan). Anterior is to the left. White arrowheads indicate discontinuities in the apical actin ring of the somitic epithelium, suggesting a local opening of the epithelial sheet and potential interfaces for the recruitment of additional mesenchymal cells from the somitocoel for their incorporation into the existing epithelium.

**Fig S7:**
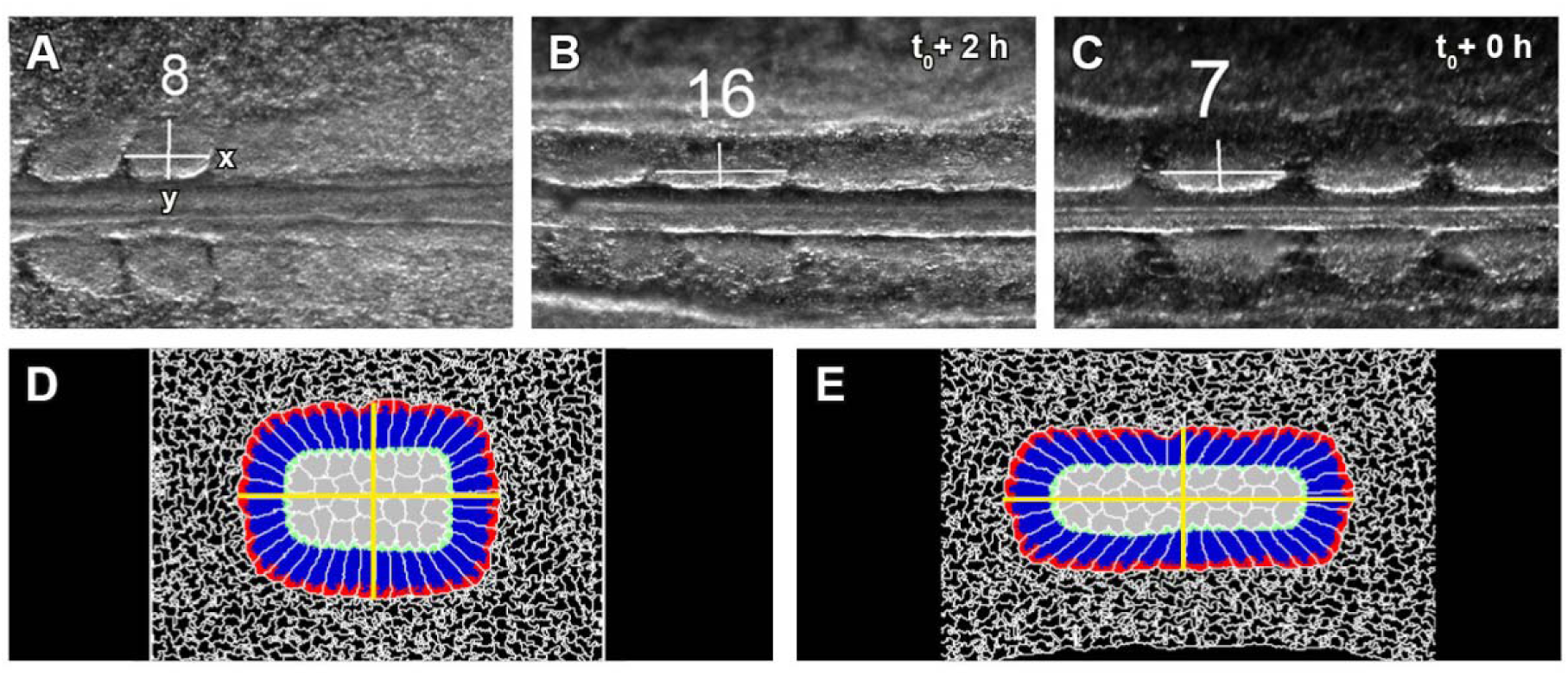
Determination of somite aspect ratios. Measuring the length (x) and width (y) of somites upon separation from the anterior tip of the PSM in control embryos (A) and stretched embryos (B), and after application of the second pull (C). Somites in (B) and (C) underwent division later. Anterior is to the left. Aspect ratio determination *in silico* before (D) and after (E) the pull.

**Fig S8:**
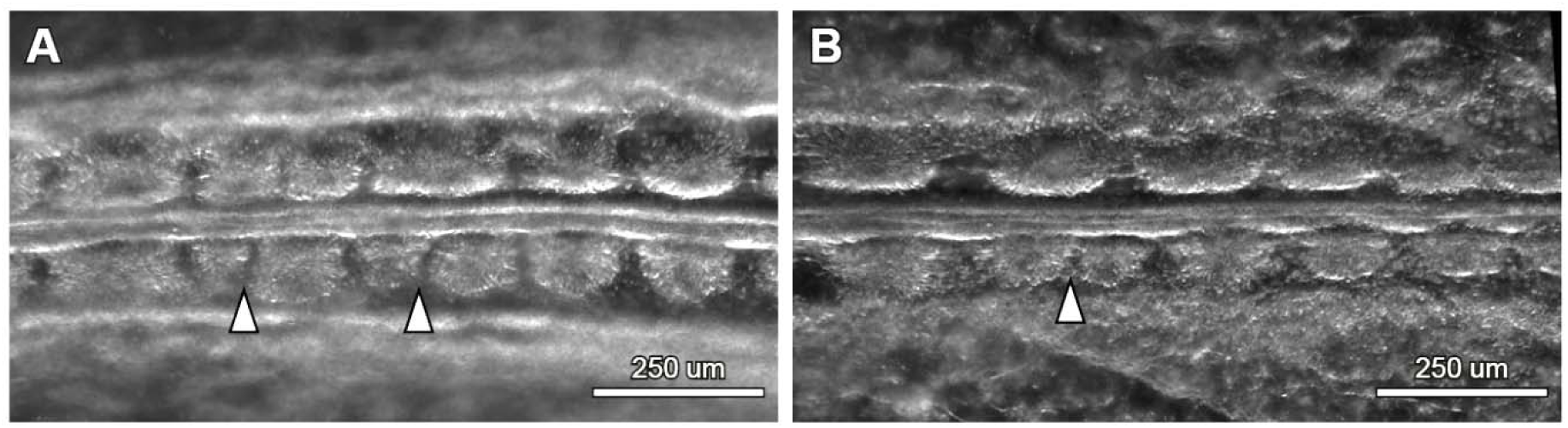
Unequally sized daughter somites. Anterior is to the left. White arrowheads indicate gaps between unequal daughter somite pairs.

**Fig S9:**
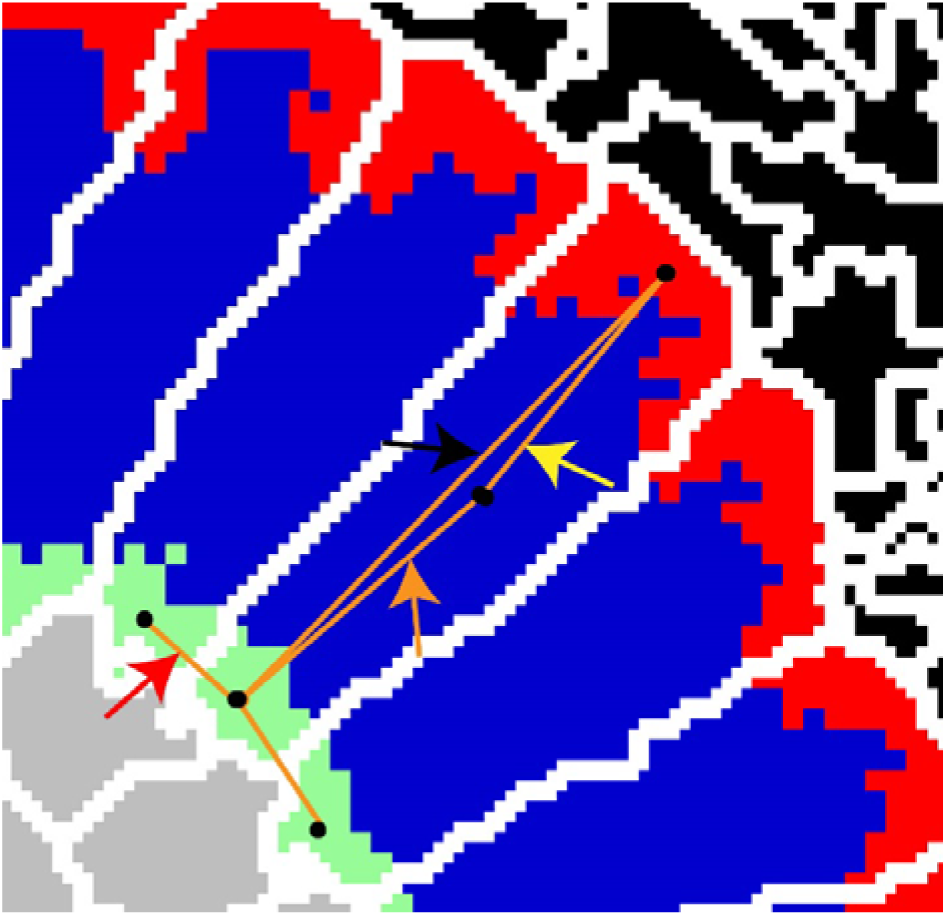
Snapshot of various cell types present in the simulations. Mesenchymal cells shown in grey. Epithelial cells consist of three domains: Apical (green), lateral (blue) and basal (red) domain. Center of masses (black dots) of epithelial cell domains are connected internally to each other using elastic springs. The springs between apical and lateral domain (orange arrow), lateral and basal domain (yellow arrow) and apical and basal domain (black arrow) are indicated. Each epithelial cell has the same spring configuration helping epithelial cells to elongate after polarization. Apical domains of neighboring are also connected via springs (red arrow) to trigger the formation of single layer of epithelial cells. Black cells in the upper right-hand corner represent ECM.

**Fig S10:**
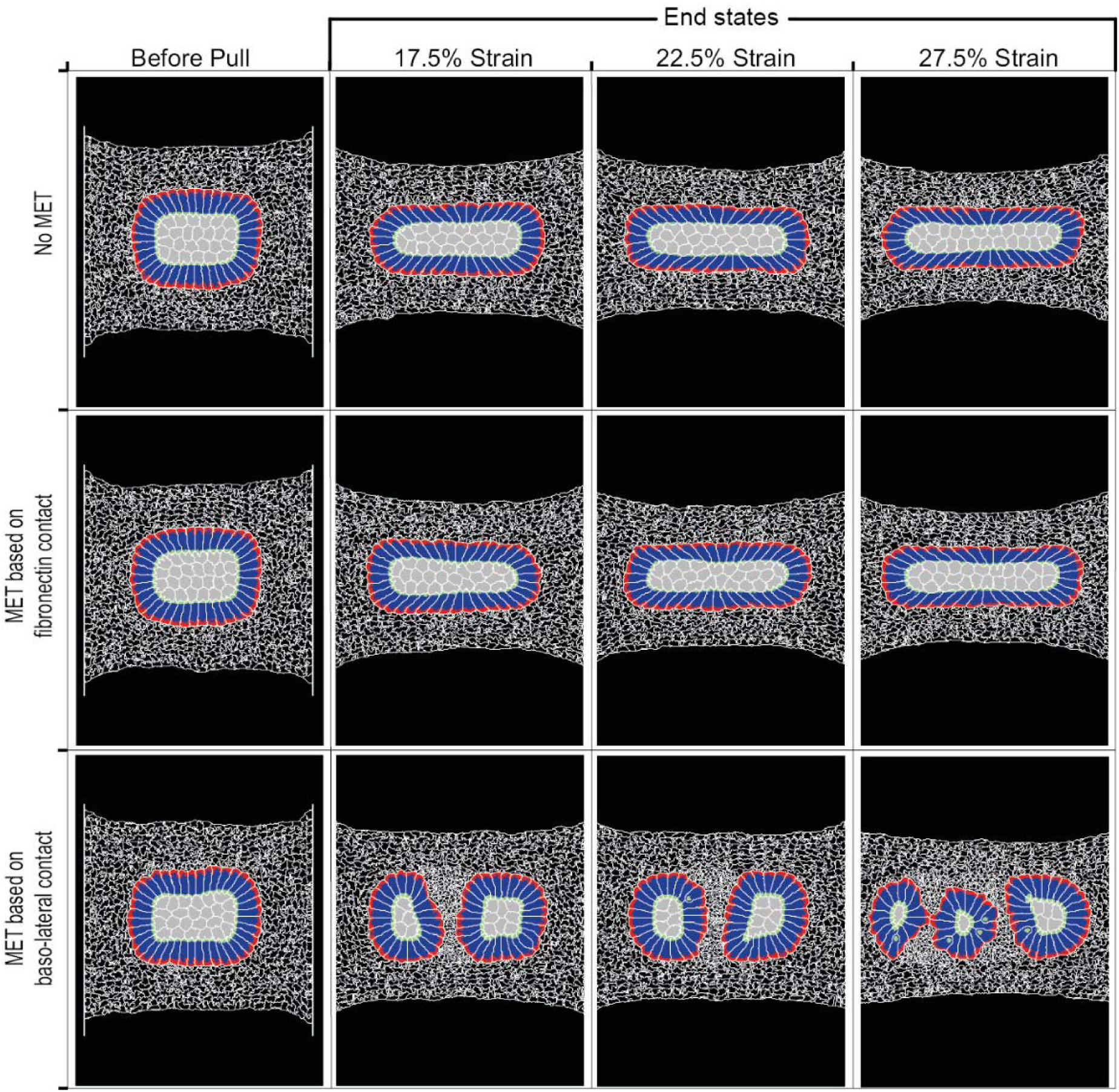
End states of simulations, testing different mesenchymal-epithelialization transition rules. Fully epithelialized somites were exposed to different strains (strain given by relative change in distance between movable walls), inducing aspect ratios of 2.5 (17.5% strain), 2.9 (22.5% strain) and 3.3 (27.5%) (compare Fig 3G). *Top row*: No MET was allowed after the pull. This was due to our initial hypothesis that somite doubling is nothing more than the reorganization of existing epithelial cells. We applied different strain values to see if high strain might lead to somite doubling. No somite division was observed. *Middle row*: MET was allowed after the pull if a cell had been in contact to the surrounding ECM matrix for a certain period of time. This MET rule did not allow somite division under various strain conditions. *Bottom row*: A new rule allows MET of mesenchymal cells upon contact to the lateral or basal membranes of epithelial cells. Formation of stable daughter somites could be observed. Both, ECM (fibronectin) induced epithelialization and basolateral contact induced epithelializing were sufficient to generate fully epithelialized somite-like structures prior to stretching.

**Fig S11:**
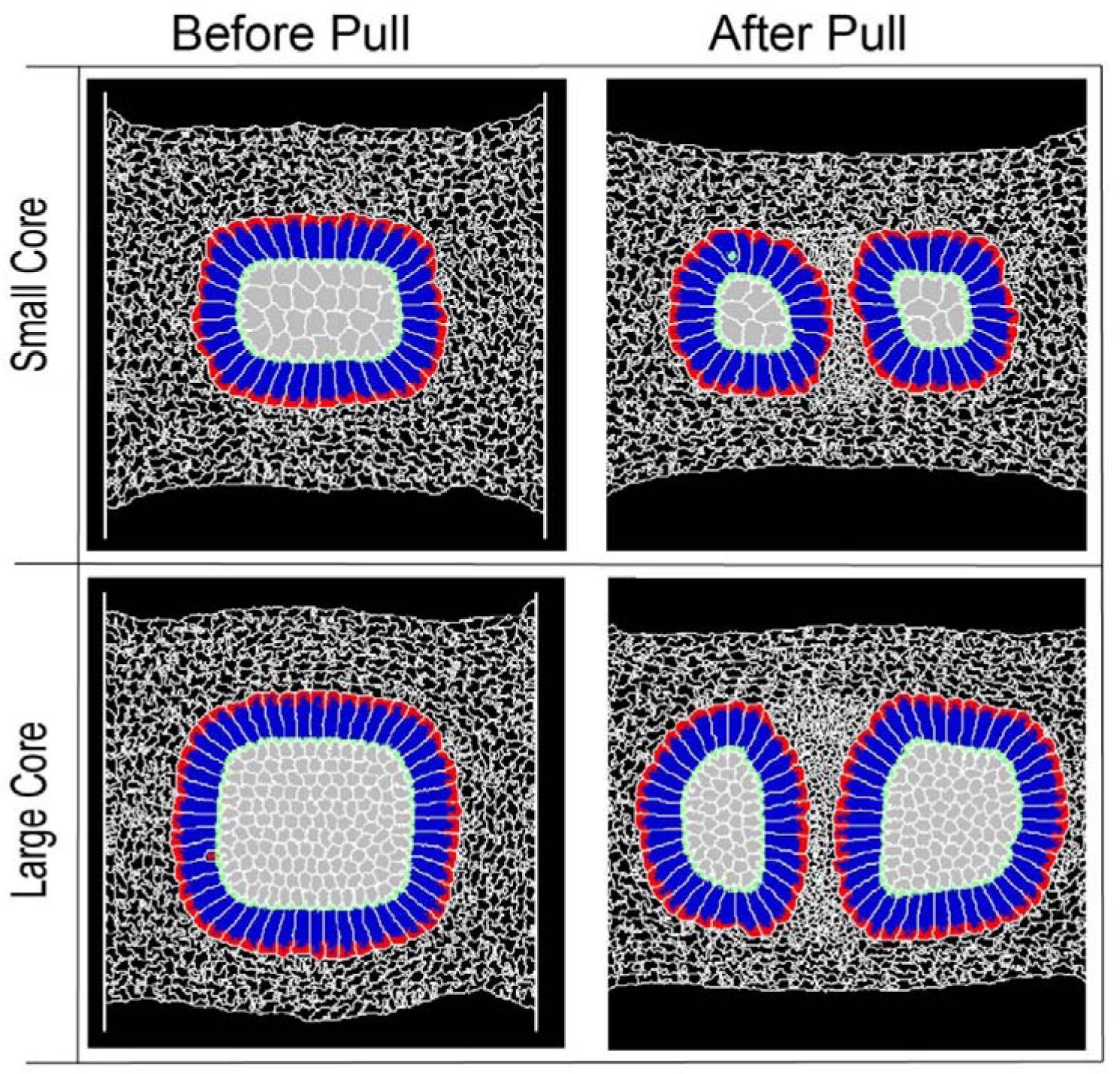
Different somite core sizes and mesenchymal cell sizes. *Top row*: Small core somite before and after pull showing somite divisions after pull. Similar response of somite division can also be seen in the large core somite (*Bottom row*). This in silico model of somite division also allows to change the size of the cells. This is evident by comparing the size of the core mesenchymal cells (in grey) in the top and bottom rows.

**Fig S12:**
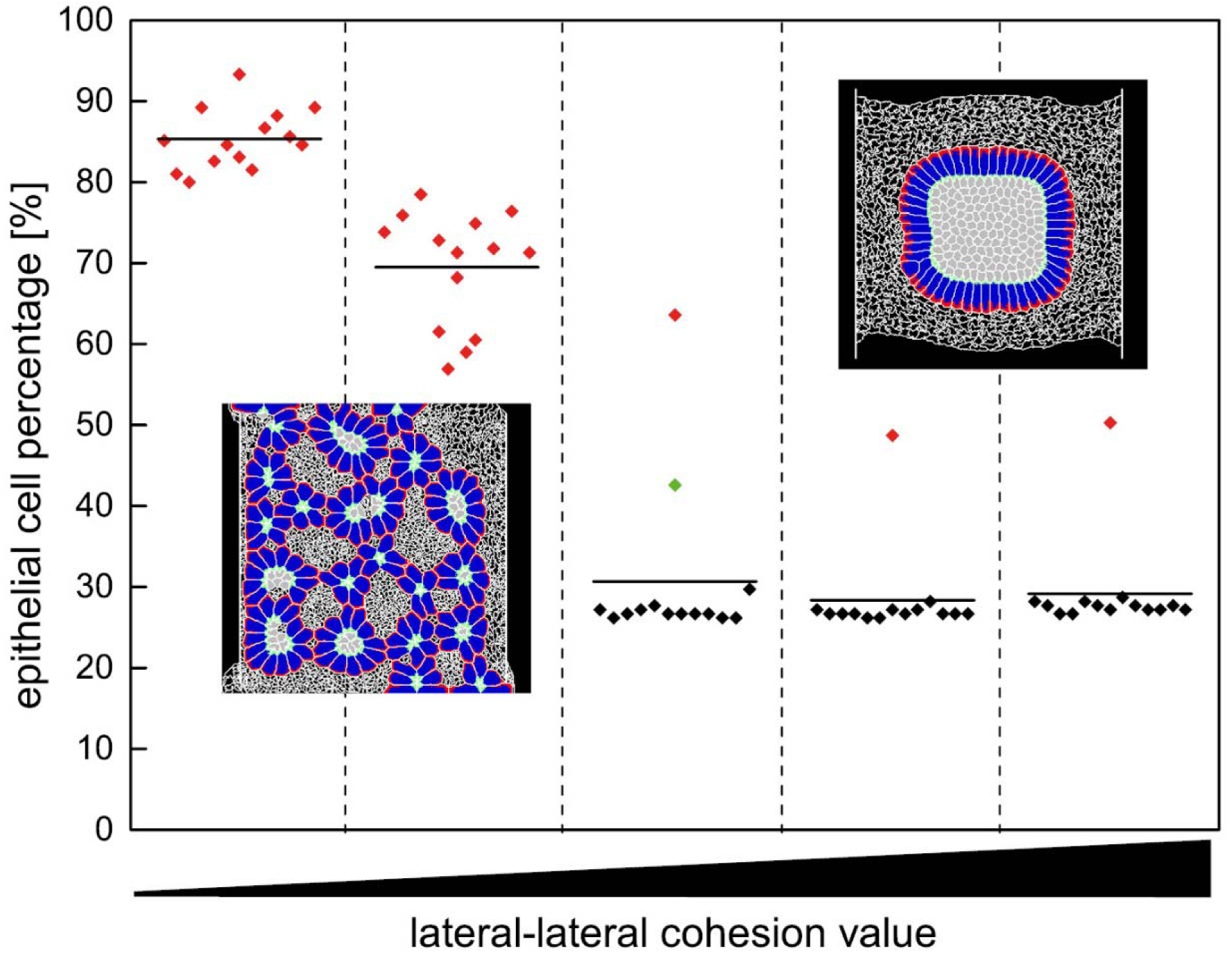
Influence of cohesion between lateral domains of epithelial cells on daughter somite number and epithelial cell percentages *in silico*. Cohesion between lateral domains of epithelial cells was varied in non-stretched somites to study the effect on somite formation. With low cohesion, almost 85% of mesenchymal cells became epithelial. This increase in epithelial cell number resulted in the formation of many epithelialized cell clusters (red data points), whereas for high cohesion values, we observed no somite division except for one case (green data point). Non-dividing somites are indicated by black data points.

**S1 Table:** **Mechanical strain caused by application of the external mechanical pull**.

| Daughter<br>somite<br>formation? | Embryo<br>No. | Distance per<br>pull<br>[mm] | Mechanical Strain<br>[%] |  |
| --- | --- | --- | --- | --- |
|  |  |  | pull 1 | pull 2 |
| Yes | 1 | 3.85 | 21 | 19 |
|  | 2 | 3.8 | 21 | 20 |
|  | 3 | 3.7 | 21 | 20 |
|  | 4 | 3.85 | 26 | 16 |
|  | 5 | 3.85 | 26 | 21 |
|  | 6 | 3.85 | 21 | 17 |
|  | 7 | 3.85 | 26 | 15 |
|  | 8 | 3.85 | 25 | 17 |
|  | 9 | 3.85 | 26 | 15 |
|  | 10 | 3.9 | 27 | 24 |
|  | 11 | 3.95 | 28 | 19 |
|  | 12 | 3.95 | 18 | 20 |
|  | 13 | 3.95 | 24 | 22 |
|  | 14 | 3.95 | 22 | 18 |
|  | 15 | 3.95 | 20 | 19 |
|  | 16 | 3.95 | 22 | 15 |
| No | 17 | 3.85 | 23 | 19 |
|  | 18 | 3.8 | 22 | 18 |
|  | 19 | 3.85 | 21 | 19 |
|  | 20 | 3.85 | 25 | 13 |
|  | 21 | 3.7 | 18 | 17 |

**S2 Table:** **Somite formation rates for stretched embryos, with and without daughter somite formation, and control embryos**.

| Daughter<br>somite<br>formation? | Embryo<br>No. | Somite No.<br>initial | Somite No.<br>final | Time<br>interval<br>[min] | Somite<br>formation<br>rate<br>[min/somite] |
| --- | --- | --- | --- | --- | --- |
| YES | 1 | 10 | 18 | 610 | 76.3 |
|  | 2 | 9 | 19 | 785 | 78.5 |
|  | 3 | 14 | 22 | 636 | 79.5 |
|  | 4 | 14 | 20 | 498 | 83.0 |
|  | 5 | 13 | 20 | 618 | 88.3 |
|  | 6 | 14 | 21 | 536 | 76.6 |
|  | 7 | 14 | 20 | 520 | 86.7 |
|  | 8 | 12 | 19 | 524 | 74.9 |
|  | 9 | 13 | 21 | 584 | 73.0 |
|  | 10 | 6 | 13 | 565 | 80.7 |
|  | 11 | 12 | 17 | 451 | 90.2 |
|  | 12 | 17 | 24 | 556 | 79.4 |
|  | 13 | 15 | 24 | 723 | 80.3 |
|  | 14 | 15 | 21 | 510 | 85.0 |
|  | 15 | 15 | 22 | 553 | 79 |
|  | 16 | 17 | 25 | 523 | 65.4 |
| Average ± Stdev.: | | | | | **79.8 ± 6.2** |
| NO | 17 | 15 | 22 | 578 | 82.6 |
|  | 18 | 11 | 21 | 795 | 79.5 |
|  | 19 | 17 | 23 | 413 | 82.6 |
|  | 20 | 15 | 21 | 510 | 85 |
|  | 21 | 12 | 22 | 826 | 82.6 |
| Average ± Stdev.: | | | | | **82.5 ± 2.0** |
| Controls | 22 | 6 | 12 | 527 | 87.8 |
|  | 23 | 6 | 23 | 1301 | 76.5 |
|  | 24 | 7 | 20 | 915 | 70.4 |
|  | 25 | 8 | 21 | 1079 | 83 |
| Average ± Stdev.: | | | | | **79.4 ± 7.6** |

**S3 Table:**
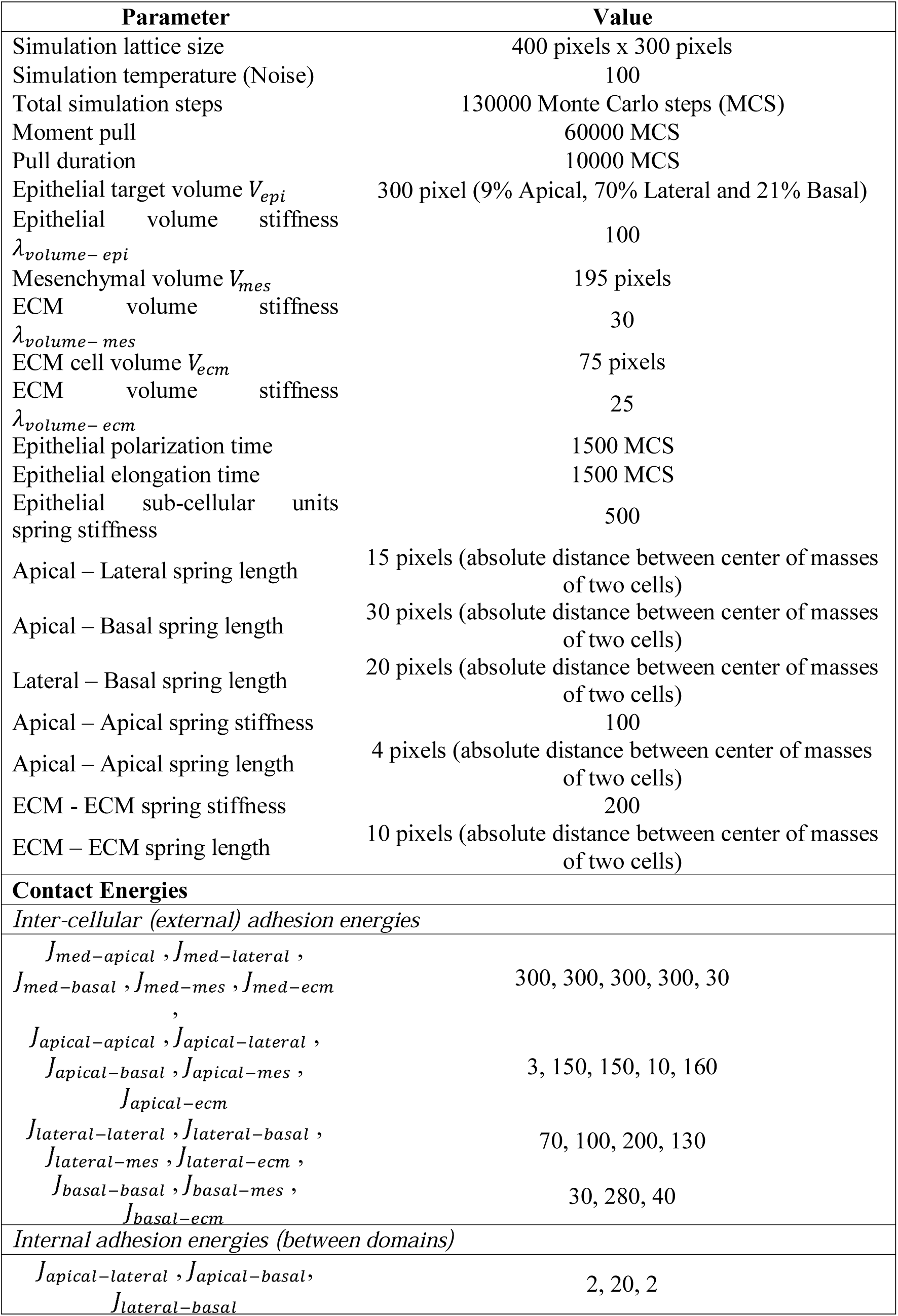
**Parameters values used in the Cellular Potts Model**.

## Movie S1: Control embryo

Time-lapse movie of the development of a control chick embryo (stage HH9+), cultured ex ovo, mounted in submerged filter paper sandwiches in the stretch setup for 19 hours, without stretching. Timer is in hours.

## Movie S2: Stretched embryo showing somite divisions

Time-lapse movie of the development of an experimental chick embryo (stage HH10), cultured ex ovo, mounted in submerged filter paper sandwiches in the stretch setup for 17,5 hours, and stretched at a speed of 1.2 μm/s along the anterior-posterior (AP) axis, in two pulls. First the overview is shown, afterwards a zoom in at the mesoderm. The stretching deforms the embryos slowly but substantially, while development progresses without damage. During the deformation, somites divide into daughter somites of different sizes, as marked by the white arrowheads in the zoom. For example: the first arrowhead shows an asymmetric somite division (lower left), while the second arrowhead shows a symmetric division (upper left).

## Movie S3: Somite epithelialization and division *in silico*

Simulation video of dividing *in silico* somites, consisting out of frames of every 500^th^ Monte Carlo Step. We tested different rules in our Cellular Potts model concerning cellular behavior in the deformed somites, to find out the most likely mechanism for the somite division to take place. Daughter somite formation in stretched somites could only successfully be induced when a MET of mesenchymal core cells upon contact to the basal or lateral membranes of epithelial cells was allowed.

